# Natural selection and the advantage of recombination

**DOI:** 10.1101/2020.08.28.271486

**Authors:** Philip J Gerrish, Benjamin Galeota-Sprung, Fernando Cordero, Paul Sniegowski, Alexandre Colato, Nicholas Hengartner, Varun Vejalla, Julien Chevallier, Bernard Ycart

## Abstract

The ubiquity of sex and recombination in nature is widely viewed as enigmatic, despite an abundance of limited-scope explanations. Natural selection, it seems, should amplify well-matched combinations of genes. Recombination would break up these well-matched combinations and should thus be suppressed. We show, to the contrary, that on average: 1) natural selection amplifies poorly-matched gene combinations and 2) creates time-averaged negative associations in the process. Recombination breaks up these poorly-matched combinations, neutralizes the negative associations, and should thus be passively and universally favored.

---

The ability to exchange genetic material through recombination (and sex) is a heritable trait [1, 2] that is influenced by many different evolutionary and ecological factors, both direct and indirect, both positive and negative. Evidence from nature clearly indicates that the net effect of these factors must be positive: recombination across all levels of organismal size and complexity is undeniably the rule rather than the exception [3–6]. Theoretical studies, on the other hand, have revealed a variety of different mechanisms and circumstances that can promote the evolution of recombination, but each one by itself is of limited scope [5–7]. These studies would thus predict that the absence of recombination is the rule and its presence an exception [8–13]. The sheer abundance of these exceptions, however, can be seen as amounting to a rule in its own right – a “pluralist” view that has been adopted by some authors to explain the ubiquity of recombination [3, 4, 14]. The necessity of this pluralist view, however, may be seen as pointing toward a fundamental shortcoming in existing theory: perhaps some very general factor that would favor recombination has been missing [3, 6, 7, 15].

Existing theories of the evolution and maintenance of sex and recombination can be divided into those that invoke *direct* vs *indirect* selection on recombination. Theories invoking direct selection propose that recombination evolved and is maintained by some physiological effect that mechanisms of recombination themselves have on survival or on replication efficiency [16–20]. Such theories might speak to the origins of sex and recombination but they falter when applied to their maintenance [21]. Most theories invoke indirect selection: they assume that any direct effect of recombination mechanisms is small compared to the trans-generational consequences of recombination.

To study how indirect selection affects recombination rate, a common approach is to model two or more fitness-related genes (or *loci*) – among which recombination may occur – as well as an additional locus, called a *modifier* locus, that determines the recombination rate. The action of natural selection on the fitness-related loci can indirectly affect the selective value and fate of different gene variants, or *alleles,* that confer different recombination rates at the modifier locus, thereby causing the mean recombination rate of the population to increase or decrease.

An allele at the modifier locus can have short-term and long-term effects [22] that can, in theory, complement or oppose each other. In the short term, modifiers that increase the recombination rate, or *up-modifiers,* will be indirectly favored if the population harbors an excess of selectively mismatched combinations of alleles across loci and a deficit of selectively matched combinations. Recombination is favored under these conditions because on average it breaks up the mismatched combinations and assembles matched combinations. Assembling selectively matched combinations increases the efficiency of natural selection: putting high-fitness alleles together can expedite their fixation [22–26], and putting low-fitness alleles together can expedite their elimination [27, 28]. In the long term, up-modifiers of recombination rate, can be indirectly favored because of the fitness variation they augment [22]. This long-term advantage was identified in the earliest speculations as to why recombination (and sex) might have evolved [29] which, curiously, predates the rediscovery of Mendel’s work and was thus written under the paradigm of Darwinian blending inheritance.

A modifier can itself be subject to the very recombination it modulates and can thus have limited-term linkage to the fitness loci whose recombination rate it modifies. Whether the selective value of recombination is determined by short-term or long-term effects depends on how long a modifier will typically remain linked to the fitness loci whose recombination rate it modifies; a loosely-linked modifier will be affected by short-term effects whereas a tightly-linked modifier will be affected by both short- and long-term effects. We derive the selective value and dynamics of a recombination-competent (*rec*^+^) modifier under loose and tight modifier linkage.

To address the evolution of sex and recombination, we have taken a reductionist approach. Our aim is restricted to studying the effects of one very key process, namely natural selection, in isolation (no mutation, no drift, etc), and we distill this problem to what we believe is its most essential form: we ask, how does the action of natural selection, by itself, affect the selective value and fate of recombination? Details of our analyses, proofs, simulation descriptions, and generalizations to *m* loci and *n* genotypes are found in companion publications ev1 [30] and ev2 [31] as well as the Supplemental Materials (SM).

We consider a large population consisting of an organism with two loci and a number of distinct alleles at each locus. An allele at the first locus contributes an amount *X* to total organismal fitness; an allele at the second locus contributes an amount *Y* to total organismal fitness; *X* and *Y* are random variables. We let *σ_XY_* denote covariance between genic fitness contributions *X* and *Y*. We find that when linkage between modifier and fitness loci is incomplete, the selective advantage of a recombination-competent (*rec*^+^) modifier in an otherwise non-recombining population is 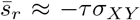, where *τ* is the characteristic duration of linkage between modifier and fitness loci. Because we are interested primarily in the sign of the selective value of recombination, we can let *τ* = 1 without loss of generality. To make our language precise, we define “loose linkage” to mean *τ* =1, and we will let 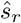 denote selective advantage under loose linkage; we note that the selective value of recombination is equal to the selective value of recombinants in this case. We define “incomplete linkage” to mean *τ* > 1, and we will let 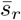 denote selective advantage under incomplete linkage.

The selective value of recombination under loose or incomplete linkage is naturally partitioned in an illuminating and biologically meaningful way by the two terms of the *total covariance*:

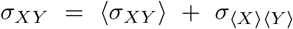

where angular brackets denote expectation in a finite population (or ensemble average). The two terms on the right-hand side correspond to the two parts of our study. When natural selection acts in isolation, the first term is transient and we are interested in its time integral as an indication of whether recombination is promoted or suppressed on average during the *process* of natural selection within local populations. The second term is not transient and we are interested in its temporal limit as an indication of whether recombination is promoted or suppressed on average between different *products* of natural selection coming from different local populations. Mathematically, the two prongs of our study thus focus on the quantities 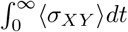 (ev2 [31]) and lim_*t*→∞_ *σ*_〈X〉〈Y〉_ (ev1 [30]).

We note that covariance is related to a commonly-used metric for selective imbalance affecting recombination, namely linkage disequilibrium (LD) [7, 22, 24, 32–35], which measures bias in allelic frequencies across loci but does not retain information about the selective value of those alleles. Covariance, on the other hand, retains information about both the frequencies and selective value of alleles. Negative LD, like negative covariance, is indicative of selective conditions that favor recombination; however, negative LD does not always favor recombination whereas negative covariance does.

To isolate the effects of natural selection, we consider large (effectively infinite) populations. For compactness of presentation, we here describe the simplest scenario in which each population consists of just two competing genotypes that differ in both of two fitness-related loci. This simple setting provides a connection to foundational evolution-of-sex studies: Fisher [36] considered the case of a single beneficial mutation arising on a variable background, thereby effectively giving rise to two competing genotypes – wildtype and beneficial mutant – that differ in both the gene with the beneficial mutation (call it the *x* gene) and its genetic background (call it the *y* gene); Muller [37] considered the case of two competing genotypes, one carrying a beneficial mutation in the *x* gene and the other in the *y* gene. Both of these approaches consider two competing genotypes that differ in both of two loci, and our qualitative findings thus apply to these foundational models and others.

Figure 1 illustrates how the simplest version of the problem is posed analytically. We consider a clonal haploid organism whose genome consists of just two fitness-related loci labeled *x* and *y*. Genetically-encoded phenotypes at these two loci are quantified by random variables *X* and *Y*, both of which are positively correlated with fitness. In each large population of such organisms, two genotypes exist: one encodes the phenotype (*X*_1_, *Y*_1_), has fitness *Z*_1_ = *ϕ*(*X*_1_, *Y*_1_) and exists at some arbitrary initial frequency *p*; the other encodes phenotype (*X*_2_, *Y*_2_), has fitness *Z*_2_ = *ϕ*(*X*_2_, *Y*_2_) and exists at initial frequency 1 – *p*. The question we ask is this: Does the action of natural selection, by itself, affect covariance between *X* and *Y* and if so, how?

**Figure 1.**
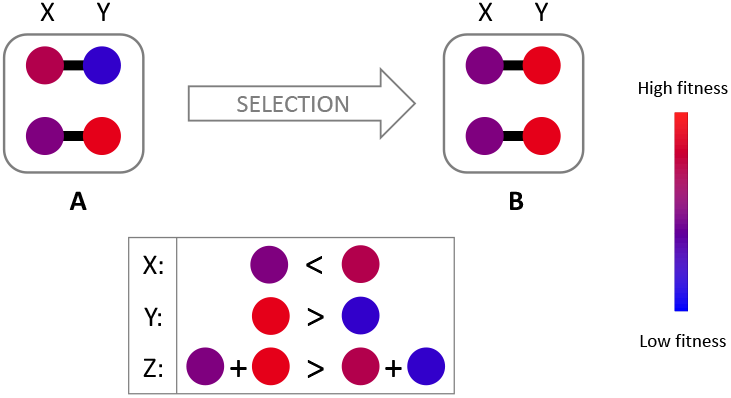
Two loci, two alleles. Here, a large (infinite) population consists of individuals whose genome has only two loci *x* and *y*, each of which carries one of two alleles: genotype 1 encodes quantified phenotype *X*_1_ at the *x* locus and *Y*_1_ at the *y* locus, and genotype 2 carries quantified phenotype *X*_2_ at the *x* locus and *Y*_2_ at the *y* locus. Fitness is indicated by color. An individual’s fitness is a function of the two phenotypes: *Z* = *ϕ*(*X,Y*); here we make the simplifying assumption that *ϕ*(*X,Y*) = *X* + *Y*, so that the fitnesses of genotypes 1 and 2 are *Z*_1_ = *X*_1_ + *Y*_1_ and *Z*_2_ = *X*_2_ + *Y*_2_, respectively. The fitter of these two genotypes has total fitness denoted *Z*^[2]^ (i.e., *Z*^[2]^ = Max{*Z*_1_,*Z*_2_}) and genic fitnesses *X*_(2)_ and *Y*_(2)_ (i.e., *Z*^[2]^ = *X*_(2)_ + *Y*_(2)_). Similarly, the less-fit of these two genotypes has total fitness *Z*^[1]^ = *X*_(1)_ + *Y*_(1)_. We note: *Z*^[2]^ > *Z*^[1]^ by definition, but this does *not* guarantee that *X*(_2_) > *X*(_1_) or that *Y*_(2)_ > *Y*_(1)_, as illustrated in the lower box. The population labeled *A* consists of two distinct genotypes but selection acts to remove the inferior genotype leaving a homogeneous population in which individuals are all genetically identical (with fitness *Z*^[2]^) as illustrated in the population labeled *B.* We derive selective mismatch measured by covariance *σ_XY_*: 1) across populations (among different *B*), given by 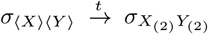, and 2) within populations (going from A to B), given by 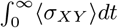.

Figure 2 illustrates the problem by analogy to a set of canoe races. On the surface, one might suspect that natural selection would promote well-matched combinations in which large values of *X* are linked to large values of *Y*, thereby creating a positive association between *X* and *Y*. In fact, this notion is so intuitive that it is considered self-evident, explicitly or implicitly, in much of the literature [3–5, 14, 21, 25, 38–41]. If this notion were true, recombination would break up good allelic combinations, on average, and should thus be selectively suppressed. Such allele shuffling has been called “genome dilution”, a label that betrays its assumed costliness. We find, however, that the foregoing intuition is flawed. To the contrary, we find that natural selection will, on average, promote an excess of mismatched combinations in which large values of *X* are linked to small values of *Y*, or vice versa, thereby creating a negative association between *X* and *Y*. Recombination will on average break up the mismatched combinations amplified by natural selection, assemble well-matched combinations, and should thus be favored.

**Figure 2.**
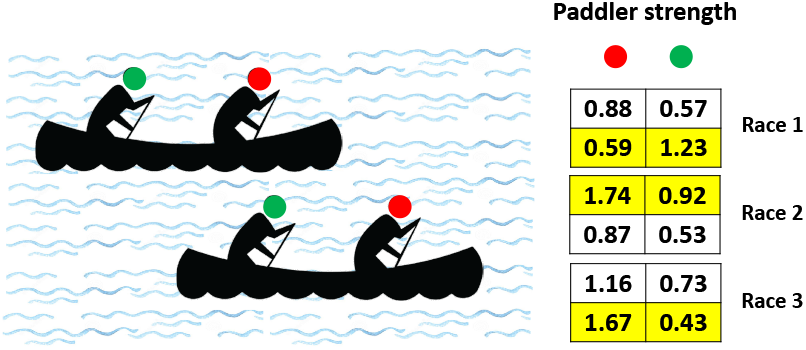
Canoe race analogy. Each canoe contains two paddlers. The strength of each paddler is measured and reported in the table. In any given canoe race, there is no correlation between paddler strengths *A* (green) and *B* (red). In each race, paddler strengths are recorded (tables on right), and the winning canoe is that in which the sum of the strengths of the two paddlers is the greatest (highlighted). Three such canoe races are conducted. We ask: what is the covariance between the strengths of paddlers *A* and *B* among winning canoes only? While it seems reasonable to suppose that winning canoes would carry two strong paddlers thereby resulting in positive covariance, the counter-intuitive answer we find is that the covariance is, for all practical purposes, unconditionally negative in expectation. By analogy, paddlers are genes, paddler strength is genic fitness, and canoes are genotypes. Natural selection picks the winner.

Figure 3 illustrates why our initial intuition was wrong and why natural selection instead tends to create negative fitness associations among genes. For simplicity of presentation, we assume here that an individual’s fitness is *Z* = *ϕ*(*X, Y*) = *X* + *Y*, i.e., that *X* and *Y* are simply additive genic fitness contributions, and that *X* and *Y* are independent. In the absence of recombination, selection does not act independently on *X* and *Y* but on their sum, *Z* = *X* + *Y*. Perhaps counter-intuitively, this fact alone creates negative associations. To illustrate, we suppose that we know the fitness of successful genotypes to be some constant, *z*, such that *X* + *Y* = *z*; here, we have the situation illustrated in Fig. 3a and we see that *X* and *Y* are negatively associated; indeed, covariance is immediate: *σ*_〈*X*〉〈*Y*〉_ = —*σ_X_σ_Y_* ≤ 0. Of course, in reality the fitnesses of successful genotypes will not be known *a priori* nor will they be equal to a constant; instead, they will follow a distribution of maxima of *Z* as illustrated in Fig. 3b. This is because, in large populations, the successful genotype will practically always be the genotype of maximum fitness. If populations consist of *n* contending genotypes, then the successful genotype will have fitness *Z*^[*n*]^ = *X*_(*n*)_ + *Y*_(*n*)_, the *n^th^* order statistic (maxima) of *Z* with genic components *X*_(*n*)_ and *Y*_(*n*)_ (called *concomitants* in the probability literature [42, 43]). In general, *Z*^[*n*]^ will have smaller variance than *Z*. Components *X*_(*n*)_ and *Y*_(*n*)_, therefore, while not exactly following a line as in Fig. 3a, will instead be constrained to a comparatively narrow distribution about that straight line, illustrated by Fig. 3b, thereby creating a negative association. Figure 3c plots ten thousand simulated populations evolving from their initial (green dots) to final (black dots) mean fitness components; this panel confirms the predicted negative association. More rigorous confirmation that selected genotypes will tend to carry selectively mismatched alleles across loci is found in the general mathematical proofs and further simulations of our companion paper (ev1 [30]).

**Figure 3.**
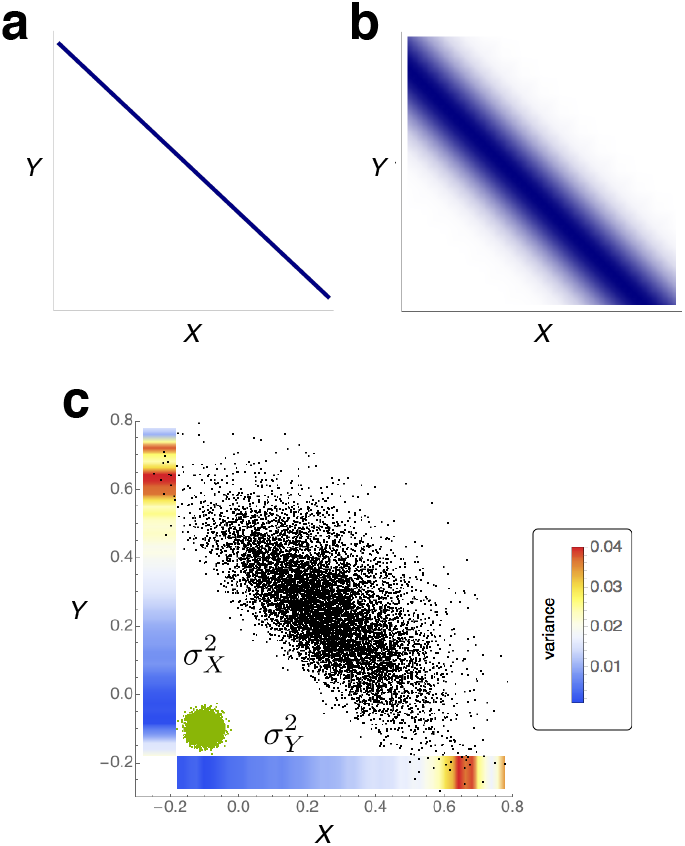
Natural selection promotes negative associations. In the absence of recombination, selection does not act independently on *X* and *Y* but organismal fitness which, for simplicity, we here assume to be their sum, *Z* = *ϕ*(*X, Y*) = *X*+*Y*. Perhaps counterintuitively, this fact alone creates negative associations. As discussed in the main text, this fact gives rise to a correlation of exactly negative one when the sum is a constant (**a**) and something intuitively negative when the sum is distributed as expected (**b**), i.e., as an order statistic. (**c**), Ten thousand simulated populations move from their initial (green dots) to final (black dots) mean fitnesses. Here, the predicted negative covariance in the final state is apparent. The heatmap bars indicate variance in *Y* along the *x*-axis and variance in *X* along the *y*-axis, a manifestation of Hill-Robertson interference [7, 28, 44–47]: larger genic fitness at one locus relaxes selection on the other locus allowing for larger fitness variance at the that locus.

What we have shown so far is that, if recombination occurs across different *products* of natural selection, the resulting offspring should be more fit than their parents, on average: lim_*t*→∞_ *ϕ*_〈*X*〉〈*Y*〉_ < 0. This effect provides novel insight into established observations that population structure can favor recombination [41, 44, 48–50] and speaks to notions that out-crossing can create hybrid vigor (heterosis) by providing a general theoretical basis for pseudo-overdominance [51–56] (explained in ev1 [30]).

Much of evolution indeed takes place in structured populations providing ample opportunity for cross-subpopulation recombination. It is thought, for example, that primordial life forms evolved primarily on surfaces that provided spatial structure [57, 58] which can enhance the advantage of recombination [5, 40, 41, 50, 59]. It is also true, however, that much of evolution takes place within unstructured (or “well-mixed”) populations; primitive life forms, for example, also existed in planktonic form [60]. We now turn to the question of how the process of evolution by natural selection affects the selective value and fate of recombination within such unstructured populations (ev2 [31]).

We begin with the case of a loosely-linked modifier. Here again, Fig. 1 shows how the problem is posed analytically. Natural selection will cause the two competing genotypes to change in frequency, causing covariance to change over time. Our measure of the net effect of natural selection on recombination under loose modifier linkage is the quantity 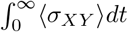; if this quantity is positive (negative), we conclude that natural selection opposes (favors) recombination on average.

In our companion paper (ev2 [31]), we show that, in expectation, time-averaged covariance is unconditionally non-positive, 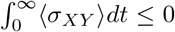, implying that *the process of natural selection always creates conditions that favor recombination, in expectation, even when modifier and fitness loci are loosely linked.*

This finding requires no assumptions about the bivariate distribution of genic fitness contributions (*X,Y*) in the initial population; in fact, a smooth density is not required (ev2 [31]). Remarkably, this distribution can even have strongly positive covariance, and yet the net effect of natural selection is still to create negative time-integrated covariance. Put differently, *recombination is advantageous, in expectation, regardless of the source of heritable variation upon which natural selection acts* – whether it be drift, migration, mutation, etc, or what the specific parameters, dynamics or interactions of these processes might be. Furthermore, our result holds unconditionally when genic fitness contributions are additive, in contrast with equilibrium-based studies that require non-additivity [22, 32, 34], and it holds when the nonadditive component (epistasis) is only loosely constrained to a wide interval about zero (ev2 [31]). Lastly, temporal fluctuations in fitness and/or epistasis, as invoked by some previous studies [32, 34, 61–64], are not required.

Our analyses further show that natural selection promotes recombination in expectation even when recombinants are present in the initial variation upon which natural selection acts, an immediate consequence of the independence of recombinant advantage on the initial fitness distribution (corroborated in ev2 [31] and SM). Put differently, even in the presence of recombination, the effect of natural selection is to promote increased recombination. The implication is that natural selection not only promotes the evolution of recombination but also its maintenance.

Until now, our focus has been on the selective value of recombination when linkage between modifier and fitness loci is incomplete (and loose). We now turn to the case of complete linkage between modifier and fitness loci.

Our analyses (ev2 [31]) show that, under complete linkage, the asymptotic selective advantage of the *rec*^+^ allele is unconditionally non-negative. This finding is again independent of the bivariate fitness distribution governing the initial variation. Our analyses further show that the expected asymptotic frequency of the *rec*^+^ allele is effectively equal to the probability that the fittest possible genotype is a virtual (or potential) recombinant. When covariance between *X* and *Y* in the initial variation is non-positive, as would be the case for example if the initial variants are themselves products of previous selection (ev1 [30]), this finding implies that the expected asymptotic modifier frequency is ≥ 1 – *n*^-(*m*-1)^, where m is the number of loci and *n* the number of alleles per locus. From this expression it is apparent that expected asymptotic modifier frequency can be very close to one under reasonable conditions. Asymptotic modifier frequency is only *not* close to one in expectation in the very unrealistic case in which the correlation coefficient of the initial fitness distribution is extremely close to +1. Remarkably, *expected asymptotic modifier frequency is independent of the strength of selection* (ev2 [31]). This observation may be seen as countering, in a sense, prevailing concerns in the literature that strong selection is required [3, 21].

Otto and Barton [5, 65] have argued that negative associations build up within a population because positive associations, in which alleles at different loci are selectively well-matched, are either removed efficiently (when they are both similarly deleterious), or fixed efficiently (when they are both similarly beneficial), thereby contributing little to overall within-population associations. Genotypes that are selectively mismatched, on the other hand, have longer sojourn times, as the less-fit loci effectively shield linked higher-fitness loci from selection. The net effect, it is argued, should be that alleles across loci will on average be selectively mismatched within a population. On one hand, our findings (ev1 [30]) diverge slightly from this *Otto & Barton effect:* we find that even genotypes that are ultimately fixed carry selectively mismatched alleles on average. On the other hand, however (ev2 [31]), our findings are entirely consistent with this effect; indeed, our proof that 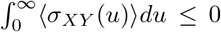, unconditionally (Prop 3 in ev2 [31]), may be interpreted as a proof of the universality of the Otto & Barton effect.

Our study identifies phenomena that are inherent consequences of natural selection and give rise to negative fitness associations across loci. Generally speaking, this pervasive phenomenon is an example of counter-intuitive effects caused by probabilistic conditioning. For example, “Berkson’s paradox” [66, 67] arises when a biased observational procedure produces spurious negative correlations. In the original context, among those admitted to hospital, it is disproportionately more likely for a patient to have, say, diabetes or, say, hypertension than to have both. Observed negative correlation between diabetes and hypertension is therefore often an artefact created by hospital admission and not representative of the general population. The selective agent in this case is hospital admission. The negative correlation may also be understood by considering that people with neither diabetes nor hypertension are less likely to be admitted to the hospital than people chosen at random from the general population.

Similarly, negative correlations arise across genic fitnesses in part because genotypes in which both loci have low genic fitness are purged by selection; in this case, however, the bias is not observational but actual, as these low-fitness genotypes are selectively removed from the population. In this genetic context, where selection is imposed artificially through breeding, this biasing effect has been contemplated and modeled; it is known as the “Bulmer effect” [68–73]. When a desired (artificially selected) trait is determined by several genetic components, one generation of breeding for the desired trait will result in negative associations among the trait’s components.

On the surface, the Bulmer effect appears similar to the phenomena we describe. On closer examination, however, it differs in several critical ways that are listed and discussed in ev1 [30] and the SM. In the context of the evolution of recombination, the most consequential difference is depicted in Fig 4. The Bulmer effect is only non-negligible under strong selection in small populations and assumes the initial distribution of fitness components is multivariate normal. In small populations, it can slightly reduce covariance from the starting covariance; if the starting covariance is positive, the Bulmer effect can reduce covariance to a slightly less-positive value, but it does not guarantee that the resulting covariance is negative. In contrast, the phenomenon we describe operates in infinite populations and results in negative time-integrated covariance independent of the initial covariance and, more generally, of the initial distribution of fitness components. In light of the above, it is clear that our findings cannot be dismissed as a simple extrapolation of the Bulmer effect.

**Figure 4.**
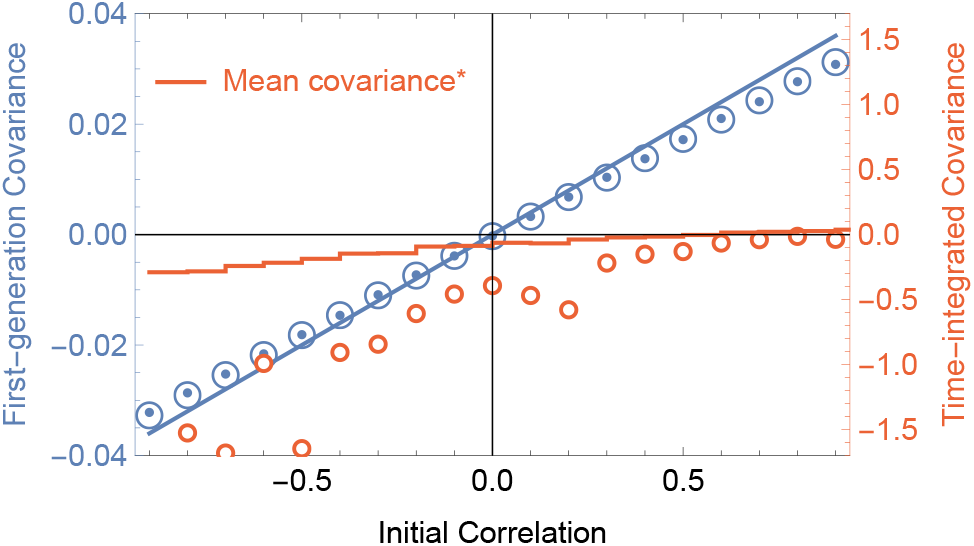
Comparison of the immediate effect (Bulmer effect) vs time-integrated effect of selection on covariance. Blue points plot simulation results after one round of selection starting with 10 genotypes whose two fitness components *X* and *Y* were pulled from a bivariate normal distribution with zero means, standard deviations *σ_X_* = 0.2 and *σ_Y_* = 0.2, and pre-selection correlation indicated by the horizontal axis and whose pre-selection covariance is the straight blue line. Open blue circles plot the theoretical expectation given by Eq S31 (SM). Our one-generation model effectively mimics the Bulmer effect although from a different theoretical starting point. Orange circles plot Montecarlo expectations of time-integrated covariance as given by Prop 3 in EV2 [31]. Orange line plots per-generation mean covariance * [multiplied by a factor of 30 to make the trend visible]. As there was no way to obtain these means analytically, plotted are the means of 500 simulations of populations of size 10^5^ initially consisting of two genotypes at equal frequency.

Our rationale for isolating natural selection was that this approach provides an in-depth understanding of one process - arguably evolution’s most influential process - by itself (an approach taken by much of classical population genetics). Most previous studies incorporate several processes simultaneously; these processes may include natural selection, mutation, drift, migration and other processes. While including several processes at once is more realistic, such an approach can make it difficult to decipher which processes are doing what. Taken together, the findings of our study clearly show that natural selection, by itself, has a remarkably encompassing tendency to create selective conditions that favor recombination, both across the *products* of selection and during the *process* of selection.

Previous studies have shown that when populations are at mutation-selection-recombination equilibrium (e.g., in the absence of adaptive evolution), decreased recombination rates are always favored, dubbed a “general reduction principle” for recombination [9, 11, 74]. Increasingly, however, empirical evidence suggests that evolution is best described as a non-equilibrium process [75–78] in which adaptive evolution is ongoing [79–83]. The setting we study is one of a population not at equilibrium. The ubiquitous recombination-augmenting tendency we describe in this paper (and companion papers ev1 [30] and ev2 [31]) may perhaps be seen as providing a “general inflation principle” of sorts, for the nonequilibrium case. Indeed, in light of the clean contrast between non-equilibrium and equilibrium findings, the ubiquity of sex and recombination in nature might be interpreted as evidence for the non-equilibrium quality of evolution generally.

Because natural selection in isolation is a transient process, the particular flavor of selection invoked (directional, stabilizing, truncation, etc) is relatively immaterial to the present study. As long as the rate of environmental change, for example, is less than the rate at which selective sorting results in fixation, our results still apply. The initial bivariate fitness distribution may reflect a population close to, at, or far from a fitness peak, for example, or it may reflect a fitness peak that is moving randomly or not.

We have found that natural selection, left to its own devices, has an encompassing tendency to promote the evolution recombination. Other more stochastic process such as mutation, migration, and drift are not explicitly modeled but are, in sense, modeled implicitly by our assumption that “something produces the heritable variation upon which natural selection acts”. In the present study, we are not concerned what that “something” is. Accounting for these other processes in the present framework is an obvious next step.

Many previous studies, in one way or another, point to the increase in agility and efficiency of adaptation that recombination confers as the primary cause of its evolution. Here, we have inverted the perspective of those earlier studies, asking not whether recombination speeds adaptation, but whether adaptation via natural selection generally creates selective conditions that promote the evolution of recombination. If so, as our findings indicate, then: 1) the ubiquity of recombination in nature might be less enigmatic than previously thought, and 2) perhaps recombination arose and is maintained more as an unavoidable byproduct than as a catalyst of natural selection.

## Acknowledgements

Much of this work was performed during a CNRS-funded visit (P.G.) to the Laboratoire Jean Kuntzmann, University of Grenoble Alpes, France, and during a visit to Bielefeld University (P.G.) funded by Deutsche Forschungsgemeinschaft (German Research Foundation, DFG) via Priority Programme SPP 1590 Probabilistic Structures in Evolution, grants BA 2469/5-2 and WA 967/4-2. P.G. and A.C. received financial support from the USA/Brazil Fulbright scholar program. P.G. and P.S. received financial support from National Aeronautics and Space Administration grant NNA15BB04A. P.G. received further support from the National Institute Of General Medical Sciences of the National Institutes of Health under Award Number R35GM137919 (awarded to Gideon Bradburd). The authors thank S. Otto and N. Barton for their thoughts on early stages of this work. Special thanks go to E. Baake for her thoughts on later stages of this work and help with key mathematical aspects. The authors thank D. Chencha, J. Streelman, R. Rosenzweig and the Biology Department at Georgia Institute of Technology for critical infrastructure and computational support.

## Footnotes

* This article is published in concert with two companion papers referenced as ev1 [30] and ev2 [31] and Supplemental Materials referenced by the adding the prefix “S”.

