## Supplementary material for "Natural selection and the advantage of recombination": SM

#### Natural selection and the advantage of recombination [S1] and companion publications [S2, S3]*

(Dated: 1 June 2022)

**A word about cross-referencing among companion publications:**

This Supplemental Materials (SM) supports three companion publications.

Cross-references among this SM and the three companion publications will appear as follows:

- Reference to the main paper (the leading paper) appears as: EV0 [#1]
- Reference to the first companion paper appears as: EV1 [#2]
- Reference to the second companion paper appears as: EV2 [#3]

where #1, #2, and #3 are the local reference numbers for the companion papers, as listed in the local bibliography.

References to objects in the SM will appear as: S# where # is the number of the referenced object.

* These Supplemental Materials support three companion papers published in concert; they are referenced as EV0 [S1] , EV1 [S3] and EV2 [S2].

†

### S1. COVARIANCE, THE MEASURE OF RECOMBINANT ADVANTAGE

**Most relevant to:** EV0 [S1] , EV1 [S3] , and EV2 [S2]

#### A. Classical discrete-time definitions

Using standard notation and definitions, we let $w_i$ denote the mean number of offspring that the $i^{th}$ individual contributes to the next generation, and $\bar{w} = \frac{1}{N}\sum_{i=1}^{N} w_i$, the grand mean number of offspring taken across all individuals in the population. The relative fitness of the $i^{th}$ individual is simply:

$$\frac{w_i}{\bar{w}} \tag{S1}$$

We emphasize here that this definition of relative fitness is in *discrete time*, where the discrete time interval is a generation. While it makes biological sense and is the most commonly-used definition of relative fitness, technically speaking it implicitly makes the questionable assumption of synchronous reproduction, which can introduce error.

We now suppose that each individual has two genes, having genic fitnesses $x$ and $y$. In the absence of epistasis, the relative fitness of the $i^{th}$ individual is:

$$\frac{x_i y_i}{\overline{xy}} \tag{S2}$$

If a recombinant is produced that carries the $x$ allele from the $i^{th}$ individual and the $y$ allele from the $j^{th}$ individual, the relative fitness of the recombinant will be:

$$\frac{x_i y_j}{\overline{xy}} \tag{S3}$$

If a recombinant is produced that carries the $x$ allele from the $i^{th}$ individual and the $y$ allele from a randomly chosen individual in the population, then on average, the relative fitness of the recombinant will be:

$$\frac{x_i \frac{1}{N}\sum_{j=1}^{N} y_j}{\overline{xy}} \tag{S4}$$

Following the same logic, if a recombinant is produced from two randomly-chosen parents, then on average, the relative fitness of the recombinant will be:

$$\bar{w}_r = \frac{\frac{1}{N}\sum_{i=1}^{N} x_i \frac{1}{N}\sum_{j=1}^{N} y_j}{\overline{xy}} \quad \xrightarrow{N\to\infty} \quad \frac{\mathbb{E}[x]\mathbb{E}[y]}{\mathbb{E}[xy]} \tag{S5}$$

The standard discrete-time definition of selective advantage is mean relative fitness after one generation minus one. By this definition, the mean selective advantage of recombinants is:

$$\bar{s}_r = \bar{w}_r - 1 = \frac{\mathbb{E}[x]\mathbb{E}[y]}{\mathbb{E}[xy]} - 1 = \frac{-\texttt{Cov}(x,y)}{\bar{w}} \tag{S6}$$

where $\bar{w} = \mathbb{E}[xy]$, the mean fitness of the population.

##### B. Continuous time

Elsewhere, we have shown that time-integrated recombinant selective advantage evolves as:

$$\int_0^t s_r(u)du \;=\; \mathcal{C}_0(t,0)+\mathcal{C}_0(0,t)-\mathcal{C}_0(t,t)$$

where $\mathcal{C}_0(\varphi,\theta)$ is the *cgf* of the initial fitness distribution.

Average recombinant advantage over the course of its first $\tau$ generation of growth is therefore:

$$\begin{aligned}\hat{s}_r(t) &= \frac{1}{\tau}\int_t^{t+\tau} s_r(u)du \\ &= \frac{1}{\tau}\left[\mathcal{C}_t(\tau,0)+\mathcal{C}_t(0,\tau)-\mathcal{C}_t(\tau,\tau)\right] \\ &= \frac{1}{\tau}\ln\frac{\mathbb{E}_t[e^{\tau X}]\mathbb{E}_t[e^{\tau Y}]}{\mathbb{E}_t[e^{\tau(X+Y)}]} \;\approx\; -\tau\sigma_{XY}(t)\end{aligned}$$

The approximation made by the last step is extremely accurate for small $\tau$ (see Figs S1 and S2) because the two-dimensional Jensen gaps for numerator and denominator essentially cancel each other out:

> Jensen's inequality in two dimensions gives $\ln\mathbb{E}[e^X]\mathbb{E}[e^Y]=\mathbb{E}[X]\mathbb{E}[Y]+JG1$ and $\ln\mathbb{E}[e^{X+Y}]=\mathbb{E}[XY]+JG2$, where $JG1$ and $JG2$ are Jensen gaps one and two. If $JG1\approx JG2$ then $\ln\mathbb{E}[e^X]\mathbb{E}[e^Y]-\ln\mathbb{E}[e^{X+Y}]\;\approx\;\mathbb{E}[X]\mathbb{E}[Y]-\mathbb{E}[XY]\;=\;-\sigma_{XY}$. Figure S1 reveals that $JG1$ and $JG2$ are indeed extremely close and thus effectively cancel each other out.

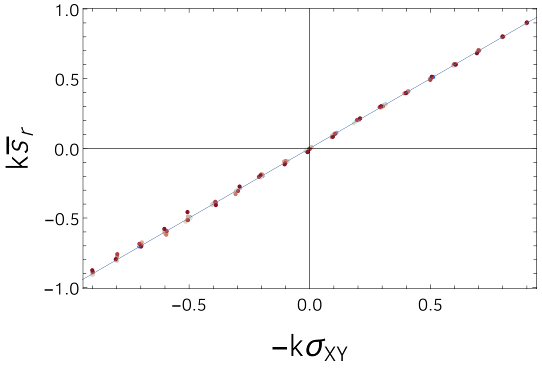


Figure S1. Mean selective advantage of recombination under loose linkage ($\tau = 1$) is extremely well approximated by minus the covariance ($-\sigma_{XY}$). Dots plot values based on 10,000 points drawn from a bivariate normal distribution with zero means and standard deviations varying between zero and one. Quantities were standardized to fit on the same plot by multiplying by $k = (\sigma_X\sigma_Y)^{-1}$ (i.e., by converting to correlation). The thin line plots the $y = x$ line.

#### S2. SIMULATIONS OF ACROSS-POPULATION COVARIANCE

**Most relevant to:** EV0 [S1] and EV1 [S3]

Here we give some Mathematica code for the simple simulations described in section III of EV1 [S3]. Figure S6 plots the across-population covariance computed by this code for different parent distributions.

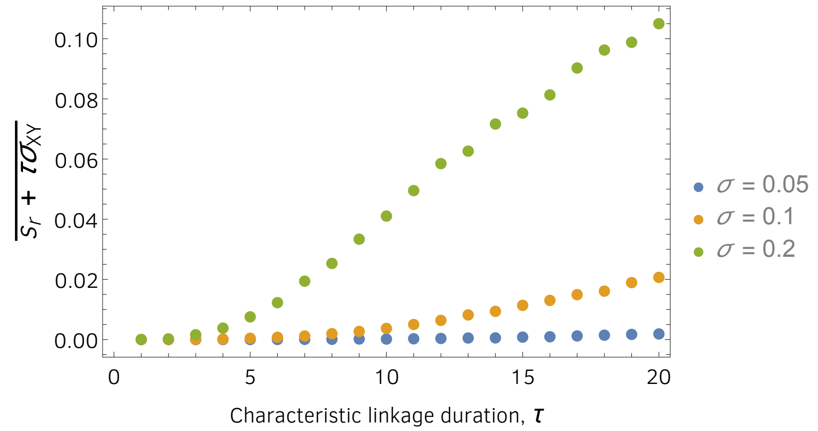


Figure S2. Mean difference between selective advantage of recombination under incomplete linkage, $\bar{s}_r = \tau^{-1} \ln \mathbb{E}[\tau X]\mathbb{E}[\tau Y]/\mathbb{E}[\tau(X+Y)]$, and its approximation $-\tau\sigma_{XY}$. Dots plot values based on 10,000 points drawn from a bivariate normal distribution with zero means and standard deviations specified in the legend. As $\tau$ increases, the difference becomes increasingly positive, indicating that the approximation is conservative for large $\tau$: $\bar{s}_r \geq -\tau\sigma_{XY}$

```
In[1]:= ng = 20; (* number of distinct genotypes in a subpopulation *)
        cr = .05; (* corr(X,Y) in initial population *)
        dist = BinormalDistribution[{-.1, -.1}, {.1, .1}, cr];
        rm = Table[
           MaximalBy[RandomReal[dist, ng], Total][[1]], {1000}];
        Covariance[rm][[1]][[2]]

Out[4]= -0.00334063
```

Figure S3. Mathematica code for simplest simulations of across-population covariance. Here, a bivariate normal distribution is used to generate $n = 20$ genic fitness pairs $((x_i, y_i),\ i = 1, 2, ..., n)$. Parameter `cr` is the correlation coefficient for these fitness pairs. The pair whose sum $x_i + y_i$ is maximum is recorded in a new array $(\hat{x}_i, \hat{y}_i)$ (denoted `rm` in the code). This is repeated 1000 times, and $\text{cov}(\hat{x}_i, \hat{y}_i)$ is reported.

```
In[1]:= ng = 20; (* number of distinct genotypes in a subpopulation *)
        d = GammaDistribution[a, b]; (* insert any 2-parameter distribution here *)
        sol = FindRoot[{Mean[d] == .1, StandardDeviation[d] == .2}, {{a, .1}, {b, .1}}];
        dist = d /. sol;
        rm = Table[
           MaximalBy[RandomReal[dist, {ng, 2}], Total][[1]], {1000}];
        Covariance[rm][[1]][[2]]

Out[6]= -0.15223
```

Figure S4. Similar to Fig S3, but here initial genic fitnesses can be drawn from any distribution with defined mean and variance, and here correlation of the initial bivariate distribution is assumed to be zero ($X$ and $Y$ are independent).

```
In[20]:= d = StudentTDistribution[a, b, c]; (* any distribution goes here *)
params = 3;  (* number of parameters in the distribution *)
Which[params == 1,
 sol = FindRoot[Quantile[d, .5] == .1, {a, .1}];,
 params == 2,
 sol = FindRoot[{Quantile[d, .5] == .1, Quantile[d, .7] == .2}, {{a, .1}, {b, .1}}];,
 params == 3,
 sol = FindRoot[{Quantile[d, .5] == .1, Quantile[d, .7] == .2, Quantile[d, .95] == .5},
    {{a, .1}, {b, .1}, {c, .1}}];
];
dist = d /. sol;

ng = 20;  (* number of distinct genotypes in a subpopulation *)
rm = Table[
  MaximalBy[RandomReal[dist, {ng, 2}], Total][[1]], {1000}];
Covariance[rm][[1]][[2]]

Out[24]= -0.111848
```

Figure S5. A generalization of Figs S3 and S4 to distributions with up to 3 parameters and with or without defined moments. This is the code used to generate Fig S6. We rather arbitrarily assigned quantiles 0.5, 0.7 and 0.95 the values of 0.1, 0.2 and 0.5, respectively, very roughly approximating the quantiles of a normal distribution with mean and standard deviation equal to 0.1 and 0.2, respectively.

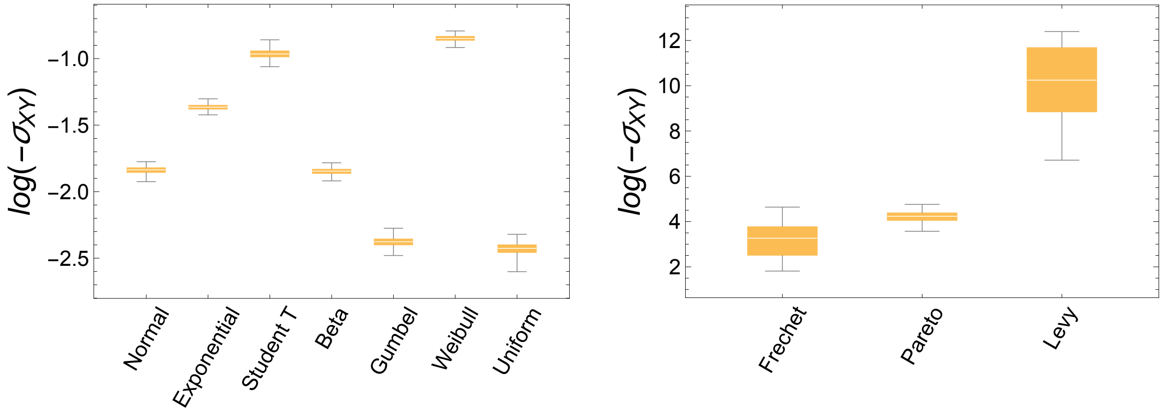


Figure S6. Across-population covariance for different parent distributions governing the initial genic fitnesses in each population. The simulation code and parameters used to generate these plots are shown in Fig S5.

#### S3. GENERAL $m$-LOCUS $n$-ALLELE SETTING

**Most relevant to:** EV0 [S1] and EV1 [S3]

##### A. Setting

Let $n$ and $m$ be two positive integers. Let $X$ be a $m$-dimensional random vector. No particular hypothesis is made at this point on the distribution of $X$: its coordinates need not be independent, nor identically distributed. For

$i = 1, \ldots, n$, let $X_i = (X_{i,j})_{1 \leqslant j \leqslant m}$ be i.i.d. copies of $X$. Let $\phi$ be a measurable function from $\mathbb{R}^m$ into $\mathbb{R}$. For $i = 1, \ldots, n$, denote by $\varphi_i$ the image by $\phi$ of the vector $X_i$.

$$\varphi_i = \phi(X_i) \ .$$

The $n$ random variables $(\varphi_i)_{1 \leqslant i \leqslant i}$ are i.i.d. Denote by $\sigma \in \mathcal{S}_n$ the random permutation such that

$$\min_{i=1}^{n} \varphi_i = \varphi_{\sigma(1)} \leqslant \cdots \leqslant \varphi_{\sigma(n)} = \max_{i=1}^{n} \varphi_i \ .$$

The permutation $\sigma$ is uniquely defined up to the usual convention of increasing order for indices corresponding to ties.

For $1 \leqslant i \leqslant n$, denote by $X_{(i)}$ the vector $X_{\sigma(i)}$, i.e. that vector whose image by $\phi$ ranks $i$-th. That notation is customary for order statistics of real-valued random variables. Here,

$$(\varphi_{(i)})_{1 \leqslant i \leqslant n} = (\varphi_{\sigma(i)})_{1 \leqslant i \leqslant n} = (\phi(X_{(i)}))_{1 \leqslant i \leqslant n}$$

are the order statistics of the vector $(\varphi_i)_{1 \leqslant i \leqslant n}$.

In the application that we have in mind, $i$ is the index of an individual in a population of size $n$, $j$ is the index of a gene contributing to the overall fitness. The random variable $X_{i,j}$ is the logarithm of the contribution of gene $j$ to the fitness of individual $i$. Thus the fitness of individual $i$ is:

$$\prod_{j=1}^{m} \mathrm{e}^{X_{i,j}} = \exp\left(\sum_{j=1}^{m} X_{i,j}\right) \ .$$

Ranking individuals by increasing fitness is ranking them by increasing values of the product of $\mathrm{e}^{X_{i,j}}$'s, or equivalently by increasing values of the sum of $X_{i,j}$'s. Thus we can take the ranking function $\phi$ to be the sum:

$$\phi(x_1, \ldots, x_m) = \sum_{i=1}^{m} x_j \ .$$

#### B. Densities

We begin with the following technical lemma, which says that the distribution of the vector which ranks $i$-th is the same as the conditional distribution of any vector given it ranks $i$-th.

Lemma S3.1. *For all $i = 1, \ldots, n$, the distribution of $X_{(i)}$ is the conditional distribution of $X_1$ given $\sigma(i) = 1$ (i.e. $\phi(X_1)$ ranks $i$-th).*

*Proof.* For any continuous bounded function $\psi$ of $m$ variables:

$$\begin{aligned} \mathbb{E}(\psi(X_{(i)})) &= \sum_{\ell=1}^{n} \frac{1}{n} \mathbb{E}(\psi(X_\ell) \,|\, \sigma(i) = \ell) \\ &= \mathbb{E}(\psi(X_1) \,|\, \sigma(i) = 1) \ . \end{aligned}$$

Thus the distribution of $X_{(i)}$ and the conditional distribution of $X_1$ given that $\phi(X_1)$ ranks $i$-th, are the same. □

As customary, boldface letters will denote real vectors: $\mathbf{x} = (x_1, \ldots, x_m)$. From now on, we assume that the distribution of $X$ has a probability density function (pdf). Denote it by $f(\mathbf{x})$. Denote by $H$ the cumulative distribution function (cdf) of $\phi(X)$:

$$H(t) = \mathbb{P}(\phi(X) \leqslant t) = \int_{\{\phi(\mathbf{x}) \leqslant t\}} f(\mathbf{x})\,\mathrm{d}\mathbf{x}\ .$$

Denote by $H_i(t)$ the cdf of $\varphi_{(i)}$. Its expression is well known.

$$H_i(t) = \sum_{k=i}^{n} \binom{n}{k} H^k(t)(1 - H(t))^{n-k}\ . \tag{S7}$$

Proposition S3.1 below gives the pdf of $X_{(i)}$.

Proposition S3.1. *Assume $H$ is continuous, i.e. the probability of ties in the ranking is null. Then $X_{(i)}$ had a pdf, denoted by $g_i$. It is related to $f$ and $H$ by:*

$$g_i(\mathbf{x}) = n\, f(\mathbf{x}) \binom{n-1}{i-1} H^{i-1}(\phi(\mathbf{x}))(1 - H(\phi(\mathbf{x})))^{n-i}\ . \tag{S8}$$

Observe that

$$\frac{1}{n}\sum_{i=1}^{n} g_i = f\ ,$$

i.e. the arithmetic mean of the densities of all $X_{(i)}$'s is the common density of all $X_i$'s. This was to be expected, since choosing at random one of the $X_{(i)}$'s is equivalent to choosing any of the $X_i$'s.

*Proof.* The pdf of $X_1$ is $f(\mathbf{x})$. The probability of the event $\sigma(i) = 1$ is $1/n$. Conditioning on $X_1 = \mathbf{x}$, the probability that $X_1$ ranks $i$-th is the probability that among $\varphi_2, \ldots, \varphi_n$, $i-1$ are below $\phi(\mathbf{x})$ and $n-i$ are above. The probability for $\varphi_\ell$ to be below $\phi(\mathbf{x})$ is $H(\phi(\mathbf{x}))$. Thus the result is a consequence of Lemma S3.1. □

#### S4. EVOLUTIONARY MODELS

**Most relevant to:** EV0 [S1] and EV2 [S2]

##### A. One-locus model with selection and mutation

Let $u_t(x)$ denote probability density in fitness $x$ at time $t$ (i.e., $\int_x u_t(x) = 1$ for all $t$) for an evolving population. Dropping the subscript $t$, we have that, under selection and mutation, $u$ evolves as:

$$\partial_t u(x) = (x - \bar{x})u(x) + U \int_\gamma u(x-\gamma)g(\gamma) - Uu(x)$$

where $U$ is genomic mutation rate, and $g(\gamma)$ is probability density for fitness changes incurred by mutation, i.e., $g(\gamma)$ is the "distribution of fitness effects" of newly-arising mutations, or DFE.

Let $M(\varphi)$ denote the moment-generating function (*mgf*) for $u(x)$, i.e., $M(\varphi) = \mathbb{E}_u[e^{\varphi X}]$ and let $G(\varphi)$ denote the *mgf* for the DFE, i.e., $G(\varphi) = \mathbb{E}[e^{\varphi x}]$. The transformed equation is:

$$\partial_t M(\varphi) = \partial_\varphi M(\varphi) - \partial_\varphi M(0)M(\varphi) + UM(\varphi)(G(\varphi) - 1).$$

Over the time interval in question (assumed to be short on evolutionary time scales), we will suppose the DFE is invariant such that $G(\varphi)$ is a constant function. We define cumulant-generating function (*cgf*) $\mathcal{C}(\varphi) = \ln M(\varphi)$; noting that $\partial_\varphi \mathcal{C}(\varphi) = (\partial_\varphi M(\varphi))/M(\varphi)$, and $\partial_t \mathcal{C}(\varphi) = (\partial_t M(\varphi))/M(\varphi)$ we find that the *cgf* evolves as:

$$\partial_t \mathcal{C}(\varphi) = \partial_\varphi \mathcal{C}(\varphi) - \partial_\varphi \mathcal{C}(0) + U(G(\varphi) - 1).$$

This equation is a variant of the transport equation and, when boundary condition $\mathcal{C}(0) = 0\ \forall\ t$ is applied, it has solution:

$$\mathcal{C}_t(\varphi) = \mathcal{C}_0(\varphi + t) - \mathcal{C}_0(t) + U \int_0^t (G(\varphi + \gamma) - G(\gamma))d\gamma \tag{S9}$$

where the subscripts are now necessary again: $\mathcal{C}_t(\varphi)$ is the *cgf* of the fitness distribution $u_t(x)$ at time $t$. We note that the evolution of a population can thus be projected into the future based only on the present fitness distribution (i.e., at $t = 0$).

##### B. Two-locus model with selection and mutation

We now suppose that there are two "genes" that determine fitness, such that total fitness is determined by their sum. Letting fitness contributions of the two genes be denoted by $x$ and $y$, respectively, the total fitness is then simply $x + y$. The extension of the previous one-dimensional *pde* is therefore immediate:

Let $u_t(x, y)$ denote probability density in fitness contributions $x$ and $y$ at time $t$ for an evolving population. Dropping the subscripts again, under selection and mutation, $u$ evolves as:

$$\begin{aligned}\partial_t u(x, y) &= (x + y - \bar{x} - \bar{y})u(x, y) \\ &\quad + U \int_{\gamma,\phi} u(x - \gamma, y - \phi)g(\gamma, \phi) - Uu(x, y)\end{aligned}$$

where again $U$ is genomic mutation rate, and $g(\gamma, \phi)$ is again the "distribution of fitness effects" of newly-arising mutations, or DFE, only now it is a bivariate distribution.

Let $M(\varphi, \theta)$ denote the *mgf* for $u(x, y)$, i.e., $M(\varphi, \theta) = \mathbb{E}_u[e^{\varphi X + \theta Y}]$ and let $G(\varphi, \theta)$ denote the *mgf* for the DFE, i.e., $G(\varphi, \theta) = \mathbb{E}_g[e^{\varphi X + \theta Y}]$. Again, over the time interval in question (assumed to be short on evolutionary time scales), we will suppose the DFE is invariant such that $G(\varphi, \theta)$ is a constant function. As before, defining *cgf* $\mathcal{C}(\varphi, \theta) = \ln M(\varphi, \theta)$, we have:

$$\begin{aligned}\partial_t \mathcal{C}(\varphi, \theta) &= \partial_\varphi \mathcal{C}(\varphi, \theta) + \partial_\theta \mathcal{C}(\varphi, \theta) - \partial_\varphi \mathcal{C}(0, 0) - \partial_\theta \mathcal{C}(0, 0) \\ &\quad + U(G(\varphi, \theta) - 1).\end{aligned} \tag{S10}$$

This equation is a two-dimensional variant of the transport equation and has more possible solution forms than the one-dimensional case, namely, solutions can be of the form: $\mathcal{C}(t + \varphi, \theta - \varphi)$, $\mathcal{C}(t + \theta, \varphi - \theta)$, or $\mathcal{C}(t + \varphi, t + \theta)$. The consistent solution is the last of these. When boundary condition $\mathcal{C}(0, 0) = 0\ \forall\ t$ is applied, it has solution:

$$\mathcal{C}_t(\varphi,\theta) = \mathcal{C}_0(\varphi+t,\theta+t) - \mathcal{C}_0(t,t) + U\int_0^t (G(\varphi+\gamma,\theta+\gamma) - G(\gamma,\gamma))d\gamma \quad \text{(S11)}$$

where the subscripts have again become necessary. We again note that the evolution of a population can thus be projected into the future based only on the present fitness distribution (i.e., at $t = 0$).

##### C. Two-locus model with selection, mutation and drift

For our first few steps, we follow the logic outlined in Ewens' book [S4], section 4.10. We use the same notation as in that reference, with the exception of his use of the variable $x$ which we replace with $q$, because we will later use $X$ as we have before, to denote the fitness contribution of one of the two genes. We define the expectation of some arbitrary function $g(\mathbf{q})$, to be:

$$\mathbb{E}[g(\mathbf{q})] = \int g(\mathbf{q}) f(\mathbf{q};\mathbf{p},t)d\mathbf{q}$$

where $f(\mathbf{q};\mathbf{p},t)$ is a probability density of a diffusion process in $\mathbf{q}$ with initial frequencies $\mathbf{p}$.

Ewens [S4] (p.154, eq.4.83) gives the rate of change of the expectation of $g(\mathbf{q})$:

$$\frac{\partial}{\partial t}\mathbb{E}[g(\mathbf{q})] = \mathbb{E}\left[\sum a_i(\mathbf{q})\frac{\partial g(\mathbf{q})}{\partial q_i} + \frac{1}{2}\sum b_i(\mathbf{q})\frac{\partial^2 g(\mathbf{q})}{\partial q_i^2} + \sum\sum c_{ij}(\mathbf{q})\frac{\partial^2 g(\mathbf{q})}{\partial q_i \partial q_j}\right] \quad \text{(S12)}$$

We now define the function $g(\mathbf{q})$ to be:

$$g(\mathbf{q}) = g(q_1, q_2, ..., q_n) = \texttt{Log}\left(\sum_{i=1}^{n} q_i e^{\varphi X_i + \theta Y_i}\right) \;=\; \tilde{\mathcal{C}}(\varphi,\theta) \quad \text{(S13)}$$

where, adhering to our previous notation, the $X_i$ and $Y_i$ are fitness contributions of the two genes in question, as before, and $q_i$ denotes the frequency of individuals with total fitness $Z_i = X_i + Y_i$.

The first term on the right-hand side of (S12) is the selection term, with:

$$a_i(\mathbf{q})dt + \mathcal{O}(\delta t) = \mathbb{E}[\delta q_i] = q_i(Z_i - \mathbb{E}[Z])$$

and the second two terms are drift terms, with:

$$b_i(\mathbf{q})dt + \mathcal{O}(\delta t) = \texttt{Var}[\delta q_i] = q_i(1-q_i)/n$$

and

$$c_{ij}(\mathbf{q})dt + \mathcal{O}(\delta t) = \texttt{Cov}[\delta q_i, \delta q_j] = -q_i q_j/n$$

Plugging these definitions into Eq. (S12), we have:

First term (selection):

$$\sum_i a_i(\mathbf{q})\frac{\partial g(\mathbf{q})}{\partial q_i} = \frac{\sum_i q_i(X_i+Y_i)e^{\varphi X_i+\theta Y_i}}{\sum_i q_i e^{\varphi X_i+\theta Y_i}} - \sum_i q_i(X_i+Y_i)$$
$$= \tilde{\mathcal{C}}_0^{(1,0)}(\varphi,\theta)+\tilde{\mathcal{C}}_0^{(0,1)}(\varphi,\theta)-\tilde{\mathcal{C}}_0^{(1,0)}(0,0)-\tilde{\mathcal{C}}_0^{(0,1)}(0,0)$$

as before, where $\tilde{\mathcal{C}}_0(\theta,\phi)=\tilde{\mathcal{C}}(\theta,\phi,0)$ is the cumulant-generating function associated with random variables $X$ and $Y$ at time zero, as defined above in (S13).

Second term (drift 1):

$$\frac{1}{2}\sum_i b_i(\mathbf{q})\frac{\partial^2 g(\mathbf{q})}{\partial q_i^2} = -\frac{1}{2}\frac{\frac{1}{n}\sum_i q_i(1-q_i)e^{2\varphi X_i+2\theta Y_i}}{\left(\sum_i q_i e^{\varphi X_i+\theta Y_i}\right)^2}$$

Third term (drift 2):

$$\sum_i\sum_{j>i} c_{ij}(\mathbf{q})\frac{\partial^2 g(\mathbf{q})}{\partial q_i \partial q_j}$$
$$= \frac{-\frac{1}{n}\sum_i\sum_{j>i} q_i q_j e^{\varphi(X_i+X_j)+\theta(Y_i+Y_j)}}{\left(\sum_i q_i e^{\varphi X_i+\theta Y_i}\right)^2}$$

The two drift terms have a common denominator so the numerators can simply be added. When added, it can be rearranged so that the sum of the two drift terms is:

$$\frac{\frac{1}{2}\left(\sum q_i e^{\varphi X_i+\theta Y_i}\right)^2-\frac{1}{2}q_i e^{2\varphi X_i+2\theta Y_i}}{\left(\sum q_i e^{\varphi X_i+\theta Y_i}\right)^2}$$
$$= \frac{1}{2n}\frac{\tilde{M}(\varphi,\theta)^2-\tilde{M}(2\varphi,2\theta)}{\tilde{M}(\varphi,\theta)^2},$$

where $\tilde{M}(\varphi,\theta)=\sum_i q_i e^{\varphi X_i+\theta Y_i}$, the moment-generating function associated with random variables $X$ and $Y$. And of course we have that $\tilde{M}(\varphi,\theta)=e^{\tilde{\mathcal{C}}(\varphi,\theta)}$, so that the drift term may be rewritten as:

$$\frac{1}{2n}\left(1-e^{\tilde{\mathcal{C}}(2\varphi,2\theta)-2\tilde{\mathcal{C}}(\varphi,\theta)}\right)$$

The CGF equation now becomes:

$$\frac{\partial}{\partial t}\mathbb{E}[g(\mathbf{q})]=\frac{\partial}{\partial t}\mathbb{E}[\tilde{\mathcal{C}}(\varphi,\theta)]=\mathbb{E}\left[\tilde{\mathcal{C}}^{(1,0)}(\varphi,\theta)+\tilde{\mathcal{C}}^{(0,1)}(\varphi,\theta)-\tilde{\mathcal{C}}^{(1,0)}(0,0)-\tilde{\mathcal{C}}^{(0,1)}(0,0)+\frac{1}{2n}\left(1-e^{\tilde{\mathcal{C}}(2\varphi,2\theta)-2\tilde{\mathcal{C}}(\varphi,\theta)}\right)\right] \tag{S14}$$

Dropping the tildes and the expectations to reduce clutter, we can now write the full evolutionary model that incorporates selection, mutation, recombination (main text) and drift:

$$\frac{\partial}{\partial t}\mathcal{C}(\varphi,\theta)=\mathcal{C}^{(1,0)}(\varphi,\theta)+\mathcal{C}^{(0,1)}(\varphi,\theta)-\mathcal{C}^{(1,0)}(0,0)-\mathcal{C}^{(0,1)}(0,0)+R(e^{\mathcal{C}(\varphi,0)+\mathcal{C}(0,\theta)-\mathcal{C}(\varphi,\theta)}-1)+\frac{1}{2n}\left(1-e^{\mathcal{C}(2\varphi,2\theta)-2\mathcal{C}(\varphi,\theta)}\right) \tag{S15}$$

where $R$ is recombination rate.

We suspect this equation can no longer be solved because of the non-local arguments in the drift term (the $2\varphi$ and $2\theta$), but we can immediately see how drift affects the covariance by taking derivatives with respect to $\varphi$ and $\theta$ and setting these equal to zero:

$$\partial_t \sigma_{XY}(t) = \kappa_{1,2}(t) + \kappa_{2,1}(t) - \frac{1}{n}\sigma_{XY}(t)$$

where $\kappa_{i,j}(\mathrm{t})$ is the $(i,j)^{th}$ joint cumulant of $X$ and $Y$ at time $t$, and we recall $\sigma_{XY}(t) = \kappa_{1,1}(t)$. This shows that drift will tend to weakly push covariance towards zero from either side.

Now writing out the full equation for covariance dynamics, with selection, mutation, drift and recombination, we have:

$$\partial_t \sigma_{XY}(t) = \kappa_{1,2}(t) + \kappa_{2,1}(t) + Um_{1,1} - (R + \frac{1}{n})\sigma_{XY}(t)$$

where $m_{1,1} = \mathbb{E}(\Delta X \Delta Y)$ and $\Delta X$ and $\Delta Y$ are changes in fitness due to mutation; i.e., $m_{1,1}$ is the first joint moment of the DFE; and $R$ is recombination rate.

In Ewen's book [S4], $n$ is interpreted as the number of alleles plus one: $n = K - 1$, where $K$ is the number of alleles. And if you start with an asexual population in which all individuals have different fitnesses, then in effect you have $K = N$ alleles, where $N$ is population size, so that $n = N - 1 \approx N$.

The foregoing developments allow us to study the effects of drift in isolation:

$$\partial_t \sigma_{XY}(t) = -\frac{1}{n}\sigma_{XY}(t) \ ,$$

giving rise to the prediction that under drift only,

$$\sigma_{XY}(t) = \sigma_{XY}(0)e^{-t/n} \quad . \tag{S16}$$

We compare these predictions with simulations in Fig **??**. Simulations were individual-based and stochastic; they started with a population that was heterogeneous in with $\sigma_{XY}(0) = -0.025$, and proceeded with no selection and no mutation. Results of the simulations support the interpretation of $n$ as $N$.

#### S5. EXPECTED TIME-INTEGRATED RECOMBINANT ADVANTAGE IS ALWAYS POSITIVE: EXTENDED DERIVATIONS AND ALTERNATIVE PROOFS

**Most relevant to:** EV0 [S1] and EV2 [S2]

As we have shown in Section S1, and more specifically in Subsection S1 B, covariance times minus one equals immediate recombinant advantage, where "immediate" is more precisely defined as the advantage that builds over the first generation of growth. Here we derive the time-integral of covariance *within* an evolving population as a measure of total recombinant advantage within that population: If time-integrated covariance is positive, this implies that on average natural selection creates conditions that oppose recombination within the evolving population. On the other hand, if time-integrated covariance is negative, this implies that natural selection creates conditions that favor recombination within the evolving population, and the more strongly negative the time-integrated covariance, the more advantageous recombinants are.

#### A. Two loci, two genotypes

##### 1. Dynamics of partitioned covariance

We let $p(t)$ and $q(t) = 1 - p(t)$ denote the frequencies of superior and inferior genotypes, respectively, in a large population at time $t$. These frequencies are functions of genic fitnesses and are thus dependent on the vector $(X_1, Y_1, X_2, Y_2)$. We define:

$$\mathbf{p}(t) := p(t|X_1, Y_1, X_2, Y_2)$$

and

$$\mathbf{q}(t) := q(t|X_1, Y_1, X_2, Y_2)$$

If the population at time zero consists of half superior and half inferior genotypes, then we know from developments elsewhere that:

$$p(t|X_1, Y_1, X_2, Y_2) = \frac{e^{Z^{[2]}t}}{e^{Z^{[1]}t} + e^{Z^{[2]}t}}$$

and

$$q(t|X_1, Y_1, X_2, Y_2) = \frac{e^{Z^{[1]}t}}{e^{Z^{[1]}t} + e^{Z^{[2]}t}}$$

Dynamics of *total covariance* are:

$$\begin{aligned} \sigma^*_{XY}(t) &= \mathbb{E}[\mathbf{p}(t)X_{(2)}Y_{(2)} + \mathbf{q}(t)X_{(1)}Y_{(1)}] - \mathbb{E}[\mathbf{p}(t)X_{(2)} + \mathbf{q}(t)X_{(1)}]\mathbb{E}[\mathbf{p}(t)Y_{(2)} + \mathbf{q}(t)Y_{(1)}] \qquad \text{(S17)} \\ &\xrightarrow{t\to\infty} \ \mathtt{Cov}(X_{(2)}, Y_{(2)}) \ = \ -\frac{1}{16}\mathbb{E}^2[Z^{[2]} - Z^{[1]}] \end{aligned}$$

as proved in EV1 [S3]. If the $(X_1, Y_1, X_2, Y_2)$ are *iid* normal[S5, S6], then:

$$\sigma^*_{XY}(t) \xrightarrow{t\to\infty} \frac{\sigma^2_X \sigma^2_Y (\beta_2 - 1)}{\sigma^2_X + \sigma^2_Y} \ \leq \ 0 \ ,$$

as developed in EV1 [S3] for the general $n$-genotype case; $\beta_n$ is defined to be the variance of maxima of a sample of size $n$ drawn from a standard normal distribution. Dynamics of *within-population* covariance are:

$$\sigma_{XY}(t) = \mathbb{E}[\mathbf{p}(t)\mathbf{q}(t)(X_{(2)} - X_{(1)})(Y_{(2)} - Y_{(1)})] \ \xrightarrow{t\to\infty} \ 0 \qquad \text{(S18)}$$

*Across-population* covariance dynamics are:

$$\begin{aligned} \sigma^a_{XY}(t) &= \mathbb{E}[(\mathbf{p}(t)X_{(2)} + \mathbf{q}(t)X_{(1)})(\mathbf{p}(t)Y_{(2)} + \mathbf{q}(t)Y_{(1)})] \qquad \text{(S19)} \\ &- \ \mathbb{E}[\mathbf{p}(t)X_{(2)} + \mathbf{q}(t)X_{(1)}]\mathbb{E}[\mathbf{p}(t)Y_{(2)} + \mathbf{q}(t)Y_{(1)}] \qquad \text{(S20)} \\ &\xrightarrow{t\to\infty} \ \mathtt{Cov}(X_{(2)}, Y_{(2)}) \ = \ -\frac{1}{16}\mathbb{E}^2[Z^{[2]} - Z^{[1]}] \end{aligned}$$

If the $(X_1, Y_1, X_2, Y_2)$ are *iid* normal, then:

$$\sigma^a_{XY}(t),\ \sigma^*_{XY}(t) \xrightarrow{t\to\infty} \frac{\sigma_X^2\sigma_Y^2(\beta_2-1)}{\sigma_X^2+\sigma_Y^2} \ \le\ 0\ .$$

Proposition S5.1. *Within-population covariance integrated over time is:*

$$\begin{aligned}\int_0^\infty \sigma_{XY}(t)dt &= \int_0^\infty \mathbb{E}[\mathbf{p}(t)\mathbf{q}(t)(X_{(2)}-X_{(1)})(Y_{(2)}-Y_{(1)})]dt \\ &= (1-p)\mathbb{E}[\frac{(X_{(2)}-X_{(1)})(Y_{(2)}-Y_{(1)})}{(Z^{[2]}-Z^{[1]})}] \\ &= (1-p)\mathbb{E}[\frac{(X_2-X_1)(Y_2-Y_1)}{|Z_2-Z_1|}] \end{aligned} \tag{S21}$$

*where* $p$ *is the initial frequency of the superior genotype.*

We note that the integrand, within-population covariance, is immediate (it can be written down from first principles or derived from our *pde* approach) and does not rely on any assumptions about the parent distribution from which fitnesses are drawn.

*Proof*: We let $p$ denote initial frequency of the superior of the two genotypes, and we let $q = 1-p$ denote initial frequency of the inferior genotype. The dynamic equation for within-population covariance may be written out as:

$$\sigma_{XY}(t) = \frac{pq\mathrm{e}^{\left(Z^{[1]}+Z^{[2]}\right)t}}{\left(p\mathrm{e}^{Z^{[2]}t}+q\mathrm{e}^{Z^{[1]}t}\right)^2}(X_{(2)}-X_{(1)})(Y_{(2)}-Y_{(1)})$$

and the time-integral of covariance is:

$$\int_0^\infty \sigma_{XY}(t)dt = (X_{(2)}-X_{(1)})(Y_{(2)}-Y_{(1)})\int_0^\infty \frac{pq\mathrm{e}^{\left(Z^{[1]}+Z^{[2]}\right)t}}{\left(p\mathrm{e}^{Z^{[2]}t}+q\mathrm{e}^{Z^{[1]}t}\right)^2}dt$$

We focus our attention on the integral in the right-hand side. We will show that:

$$\int_0^\infty \frac{pq\mathrm{e}^{\left(Z^{[1]}+Z^{[2]}\right)t}}{\left(p\mathrm{e}^{Z^{[2]}t}+q\mathrm{e}^{Z^{[1]}t}\right)^2}dt \ =\ \frac{q}{Z^{[2]}-Z^{[1]}}\ ,$$

hence giving Eq (S21). We can expand the integrand as follows:

$$\frac{pq\mathrm{e}^{(Z^{[1]}+Z^{[2]})t}}{(p\mathrm{e}^{Z^{[2]}t}+q\mathrm{e}^{Z^{[1]}t})^2} = \frac{q}{Z^{[2]}-Z^{[1]}}\left(\frac{(pZ^{[2]}\mathrm{e}^{Z^{[2]}t}+qZ^{[1]}\mathrm{e}^{Z^{[1]}t})\mathrm{e}^{Z^{[1]}t}}{(p\mathrm{e}^{Z^{[2]}t}+q\mathrm{e}^{Z^{[1]}t})^2} - \frac{Z^{[1]}\mathrm{e}^{Z^{[1]}t}}{p\mathrm{e}^{Z^{[2]}t}+q\mathrm{e}^{Z^{[1]}t}}\right) \tag{S22}$$

We now define:

$$f := \mathrm{e}^{Z^{[1]}t} \quad \text{and} \quad g := \frac{-1}{p\mathrm{e}^{Z^{[2]}t}+q\mathrm{e}^{Z^{[1]}t}}$$

so that:

$$f' = Z^{[1]}\mathrm{e}^{Z^{[1]}t} \quad \text{and} \quad g' = \frac{pZ^{[2]}\mathrm{e}^{Z^{[2]}t}+qZ^{[1]}\mathrm{e}^{Z^{[1]}t}}{(p\mathrm{e}^{Z^{[2]}t}+q\mathrm{e}^{Z^{[1]}t})^2}$$

where the prime indicates derivative with respect to $t$. We note that the terms inside the parentheses in Eq (S22) are $fg'$ and $f'g$. We know from the product rule (integration by parts) that

$$\int fg' + \int f'g = fg$$

so that the integral over time of everything inside the parentheses in Eq (S22) reduces to:

$$fg \;=\; \left.\frac{-\mathrm{e}^{Z^{[1]}t}}{p\mathrm{e}^{Z^{[2]}t} + q\mathrm{e}^{Z^{[1]}t}}\right|_{t=0}^{\infty} \;=\; 0 - (-1) \;=\; 1$$

Hence the result:

$$\int_0^\infty \frac{pq\mathrm{e}^{\left(Z^{[1]}+Z^{[2]}\right)t}}{\left(p\mathrm{e}^{Z^{[2]}t} + q\mathrm{e}^{Z^{[1]}t}\right)^2} dt \;=\; \frac{q}{Z^{[2]} - Z^{[1]}} \;,$$

so that:

$$\mathbb{E}[\int_0^\infty \sigma_{XY}(t)dt] \;=\; q\mathbb{E}[\frac{(X_{(2)} - X_{(1)})(Y_{(2)} - Y_{(1)})}{Z^{[2]} - Z^{[1]}}]$$

where $q$ in Prop S5.1 is written as $1 - p$.

We observe that

$$(X_{(2)} - X_{(1)})(Y_{(2)} - Y_{(1)}) = (X_{(1)} - X_{(2)})(Y_{(1)} - Y_{(2)}) = (X_2 - X_1)(Y_2 - Y_1)$$

and that

$$Z^{[2]} - Z^{[1]} = |Z_2 - Z_1|$$

from which we have:

$$\mathbb{E}[\frac{(X_{(2)} - X_{(1)})(Y_{(2)} - Y_{(1)})}{Z^{[2]} - Z^{[1]}}] = \mathbb{E}[\frac{(X_2 - X_1)(Y_2 - Y_1)}{|Z_2 - Z_1|}]$$

□

Proposition S5.2. *We define spacings* $\Delta X = X_2 - X_1$, $\Delta Y = Y_2 - Y_1$, *and* $\Delta Z = Z_2 - Z_1 = \Delta X + \Delta Y$. *If the pairs* $(X_i, Y_i)$ *are independently drawn from a given distribution, then* $\Delta X$ *and* $\Delta Y$ *are symmetric about zero. Moreover, if* $\mathbb{E}[\sqrt{|\Delta X \Delta Y|}] < \infty$, *then*

$$\mathbb{E}[\int_0^\infty \sigma_{XY}(t)dt] \;=\; \mathbb{E}\left[\frac{\Delta X \Delta Y}{|\Delta Z|}\right] \;\leq\; 0.$$

*Proof*: The fact that $(\Delta X, \Delta Y)$ is symmetric about zero is a direct consequence of the fact that $(X_1, Y_1)$ and $(X_2, Y_2)$ are independent and have the same distribution. Note that, since $|x| + |y| \geq 2\sqrt{|xy|}$ for all $x, y \in \mathbb{R}$, we have

$$\mathbb{E}\left[\mathbb{1}_{\Delta X \Delta Y > 0} \frac{\Delta X \Delta Y}{|\Delta X + \Delta Y|}\right] \leq \frac{1}{2}\mathbb{E}[\sqrt{|\Delta X \Delta Y|}] < \infty.$$

Hence, $\mathbb{E}\left[\Delta X \Delta Y/|\Delta X+\Delta Y|\right]$ is well-defined and given by

$$\mathbb{E}\left[\frac{\Delta X\Delta Y}{|\Delta X+\Delta Y|}\right]=\mathbb{E}\left[\mathbb{1}_{\Delta X\Delta Y>0}\frac{\Delta X\Delta Y}{|\Delta X+\Delta Y|}\right]+\mathbb{E}\left[\mathbb{1}_{\Delta X\Delta Y<0}\frac{\Delta X\Delta Y}{|\Delta X+\Delta Y|}\right],$$

where, on the right-hand side, the first term is positive and finite and the second term is negative. If the latter is $-\infty$, the resut follows; if it is finite, since $(-\Delta X, \Delta Y)$ has the same distribution as $(\Delta X, \Delta Y)$, then

$$\begin{aligned}\mathbb{E}\left[\frac{\Delta X\Delta Y}{|\Delta X+\Delta Y|}\right]&=\mathbb{E}\left[\mathbb{1}_{\Delta X\Delta Y>0}\frac{\Delta X\Delta Y}{|\Delta X+\Delta Y|}\right]+\mathbb{E}\left[\mathbb{1}_{(-\Delta X)\Delta Y<0}\frac{(-\Delta X)\Delta Y}{|\Delta Y-\Delta X|}\right]\\&=\mathbb{E}\left[\mathbb{1}_{\Delta X\Delta Y>0}\Delta X\Delta Y\left(\frac{1}{|\Delta X+\Delta Y|}-\frac{1}{|\Delta Y-\Delta X|}\right)\right].\end{aligned}$$

Since for $x$ and $y$ having the same sign, we have $|x+y|>|y-x|$, the right-hand side in the previous sequence of identities is less than or equal to zero, which concludes the proof. □

Remark: The same proof works for the generalized linear fitness function $\phi(X,Y)=a+bX+cY$ with $b,c>0$. Note also that the proof tell us that $\mathbb{E}[\Delta X\Delta Y/|\Delta Z|]=0$ if and only if $\mathbb{P}(\Delta X\Delta Y=0)=1$.

Alternative proof 1

Let $f(x,y)$ denote the probability density governing random variables $\Delta X$ and $\Delta Y$. The only restrictions we place on $f(x,y)$ is that it be symmetric about zero in both $x$ and $y$ directions and that its tails be not too heavy so that the following integral exists in each quadrant of the $x$-$y$ plane. Both of these restrictions seem quite reasonable from a biological standpoint. The expectation may be thus be written as follows:

$$\mathbb{E}[\frac{\Delta X\Delta Y}{|\Delta Z|}] \;=\; \int_{-\infty}^{+\infty}\int_{-\infty}^{+\infty}\frac{xy}{|x+y|}f(x,y)dxdy$$

Symmetry about zero ($f(x,y)=f(-x,y)$ and $f(x,y)=f(x,-y)$) allows us to integrate quadrant-by-quadrant as follows:

$$\begin{aligned}\mathbb{E}[\frac{\Delta X\Delta Y}{|\Delta Z|}] &= \int_0^\infty\int_0^\infty\left(\frac{xy}{|x+y|}+\frac{(-x)y}{|(-x)+y|}+\frac{(-x)(-y)}{|(-x)+(-y)|}+\frac{x(-y)}{|x+(-y)|}\right)f(x,y)dxdy\\&= \int_0^\infty\int_0^\infty\left(\frac{2xy}{x+y}-\frac{2xy}{|x-y|}\right)f(x,y)dxdy\\&= \int_0^\infty\int_0^\infty\left(\frac{1}{x+y}-\frac{1}{|x-y|}\right)2xyf(x,y)dxdy \qquad\text{(S23)}\end{aligned}$$

We are now integrating over positive values of $x$ and $y$, for which:

$$\begin{gathered}(x+y)^2=x^2+2xy+y^2\geq x^2-2xy+y^2=(x-y)^2\\\sqrt{(x+y)^2}\geq\sqrt{(x-y)^2}\\x+y\geq|x-y|\end{gathered}$$

so that:

$$\frac{1}{x+y}-\frac{1}{|x-y|}\;\leq\;0$$

implying that Eq (S23) is non-positive, hence giving the result:

$$\mathbb{E}[\int_0^\infty \sigma_{XY}(t)dt] \ \leq \ 0$$

ALTERNATIVE PROOF 2.

Define: $X = u|X|$ and $Y = v|Y|$, where $u$, $v$ are independent $+/-1$ random variables, independent of $|X|$ and $|Y|$. Then: $|u|X| + v|Y|| = w(|X| + |Y|) + (1 - w)||X| - |Y||$, where $w = uv$. Again, $w$ is independent of $|X|$ and $|Y|$.

And:

$$\begin{aligned}\mathbb{E}[XY/|X+Y|)] &= \mathbb{E}[w|X||Y|/(w(|X|+|Y|)+(1-w)||X|-|Y||)] \\ &= \frac{1}{2}\mathbb{E}[|X||Y|/((|X|+|Y|)] - \frac{1}{2}\mathbb{E}[|X||Y|/||X|-|Y||] \\ &< 0\end{aligned}$$

since $|X| + |Y| \geq ||X| - |Y||$. □

*Proof*:

Set $U = |X|$ and $V = |Y|$; $M = \text{Max}(U, V)$, $m = \text{Min}(U, V)$. Then you can rewrite the expectation as:

$$\begin{aligned}\mathbb{E}[UV\{1/(U+V) - 1/(|U-V|)\}] &= \mathbb{E}[mM\{-2m/(M^2-m^2)\}] \\ &= -2\mathbb{E}[Mm^2/(M^2-m^2)] \leq 0\end{aligned}$$

Indeed, if the expectation is $\infty$, we get $-\infty$ as our answer. This approach removes the need to make the argument that $U + V > |U - V|$ and avoids the need to take a difference of expectations. □

#### S6. RECOMBINANT ADVANTAGE WITH EPISTASIS

**Most relevant to:** EV0 [S1] and EV2 [S2]

COROLLARY S6.1. *For any real number $\xi$, let us consider a fitness function of the form $\phi_\xi(x, y) = aX + bY + \xi g(X, Y)$, where $a, b > 0$ and $g$ is a function independent of $\xi$. Let $Z(\xi) = \phi_\xi(X_2, Y_2) - \phi_\xi(X_1, Y_1)$. Assume that for some $\varepsilon > 0$,*

$$\mathbb{E}\left[\sup_{|\xi|<\varepsilon} \frac{|\Delta X \Delta Y|}{|Z(\xi)|}\right] < \infty, \tag{S24}$$

*and that $\mathbb{P}(\Delta X \Delta Y = 0) < 1$. Then, there is $\varepsilon_0 \in (0, \varepsilon)$, such that for all $\xi \in (-\varepsilon_0, \varepsilon_0)$, we have*

$$\mathbb{E}\left[\frac{\Delta X \Delta Y}{|Z(\xi)|}\right] < 0.$$

*Proof.* Condition (S24) implies that the function $h : (-\varepsilon, \varepsilon) \to \mathbb{R}$ defined via

$$h(\xi) = \mathbb{E}\left[\frac{\Delta X \Delta Y}{|Z(\xi)|}\right]$$

is continuous. Moreover, since $\mathbb{P}(\Delta X \Delta Y = 0) < 1$, proceeding as in the proof of Proposition S5.2, we obtain that $h(0) < 0$. Hence, by continuity of $h$, we infer that there is $\varepsilon_0 \in (0, \varepsilon)$ such that $h$ is negative in $(-\varepsilon_0, \varepsilon_0)$, which concludes the proof. □

Let us now focus our attention on the fitness function $\phi_\xi(X, Y) = aX + bY + \xi XY$ with $a, b > 0$ and $\xi \in \mathbb{R}$. As before, let $Z(\xi) = \phi_\xi(X_2, Y_2) - \phi_\xi(X_1, Y_1) = (a + \xi Y_1)\Delta X + (b + \xi X_2)\Delta Y$. The case where the random variables $(|\Delta X \Delta Y|/|Z(\xi)|)_{\xi \in (-\varepsilon, \varepsilon)}$ are uniformly integrable (i.e. condition (S24) is satisfied) is covered already by Corollary S6.1. As a counterpart, the next result considers the case where the expectation of $|\Delta X \Delta Y|/|Z(\xi)|$ is infinite, and provides a simple condition to assure that the expectation of $\Delta X \Delta Y/|Z(\xi)|$ is negative (in fact, equal to $-\infty$).

Corollary S6.2. *Assume that the distribution of $(X_i, Y_i)$ has finite support, i.e. there is $K > 0$ such that $\mathbb{P}(X_i \in [-K, K], Y_i \in [-K, K]) = 1$ and that $|\xi| < (a \wedge b)/K$, where $a \wedge b$ denotes the minimum between $a$ and $b$. If we have*

$$\mathbb{E}\left[\frac{|\Delta X \Delta Y|}{|Z(\xi)|}\right] = \infty, \tag{S25}$$

*then*

$$\mathbb{E}\left[\frac{\Delta X \Delta Y}{|Z(\xi)|}\right] = -\infty.$$

*Proof*: Note first that, if $|\xi| < (a \wedge b)/K$, then $\mathbb{P}(a + \xi Y_1 \geq a - |\xi K|,\, b + \xi X_2 \geq b - |\xi K|) = 1$, and hence

$$\begin{aligned}
\mathbb{E}\left[\frac{\Delta X \Delta Y \mathbb{1}_{\Delta X \Delta Y > 0}}{|Z(\xi)|}\right] &= E\left[\frac{\Delta X \Delta Y \mathbb{1}_{\Delta X \Delta Y > 0}}{|(a + \xi Y_1)\Delta X + (b + \xi X_2)\Delta Y|}\right] \\
&= E\left[\frac{\Delta X \Delta Y \mathbb{1}_{\Delta X \Delta Y > 0}}{|(a + \xi Y_1)\Delta X| + |(b + \xi X_2)\Delta Y|}\right] \\
&\leq \frac{2}{(a \wedge b) - |\xi K|} E\left[\frac{|\Delta X \Delta Y|}{|\Delta X| + |\Delta Y|}\right] \\
&\leq \frac{1}{(a \wedge b) - |\xi K|} E\left[\sqrt{|\Delta X \Delta Y|}\right] \leq \frac{K}{a \wedge b} < \infty.
\end{aligned}$$

Therefore, condition (S25) implies that

$$\mathbb{E}\left[\frac{|\Delta X \Delta Y| \mathbb{1}_{\Delta X \Delta Y < 0}}{|Z(\xi)|}\right] = \infty,$$

and thus,

$$\mathbb{E}\left[\frac{\Delta X \Delta Y}{|Z(\xi)|}\right] = \mathbb{E}\left[\frac{\Delta X \Delta Y \mathbb{1}_{\Delta X \Delta Y > 0}}{|Z(\xi)|}\right] - \mathbb{E}\left[\frac{|\Delta X \Delta Y| \mathbb{1}_{\Delta X \Delta Y < 0}}{|Z(\xi)|}\right] = -\infty,$$

achieving the proof. □

#### S7. ASYMPTOTIC MODIFIER FREQUENCY WITH EPISTASIS

**Most relevant to:** EV0 [S1] and EV2 [S2]

Here, we assess the sensitivity to epistasis of our theoretical predictions for asymptotic modifier frequency. To this end, we performed stochastic simulations with epistasis and compared the final modifier frequency with the asymptotic predictions given by Eqs (32) and (33). Results of these simulations are plotted in Fig S7. This figure shows that asymptotic modifier frequency is little-affected by epistasis.

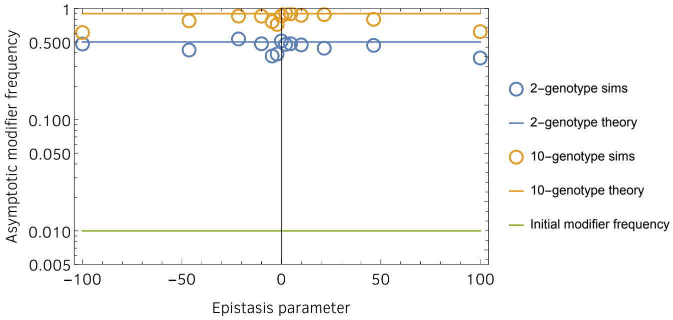


Figure S7. Effect of epistasis on asymptotic modifier frequency. Open circles plot means of 500 stochastic simulations. Blue and orange lines plot theoretical predictions given by Eq (32) for 2 and 10 genotypes, respectively. Simulations had population size 5000.

#### S8. NUMERICAL COMPUTATION OF EXPECTED TIME-INTEGRATED COVARIANCE

Recalling the definition of the empirical cumulant-generating function (*ecgf*):

$$\tilde{\mathcal{C}}_t(\varphi,\theta) \;=\; \ln \frac{1}{n}\sum_{i=1}^{n} e^{\varphi x_i+\theta y_i}$$

we have covariance dynamics:

$$\sigma_{XY}(t) \;=\; \tilde{\mathcal{C}}_0^{(1,1)}(t,t)$$

To find expected covariance, we replace the $x_i$ and $y_i$ with random variables $X_i$ and $Y_i$, and compute Montecarlo expectation by drawing these random variables at random from some bivariate continuous distribution. With this approach, we compute and plot in Figs S8 and S9 the quantity:

$$\mathbb{E}[\int_0^\infty \sigma_{XY}(u)du] \;=\; \mathbb{E}[\int_0^\infty \tilde{\mathcal{C}}_0^{(1,1)}(u,u)du]$$

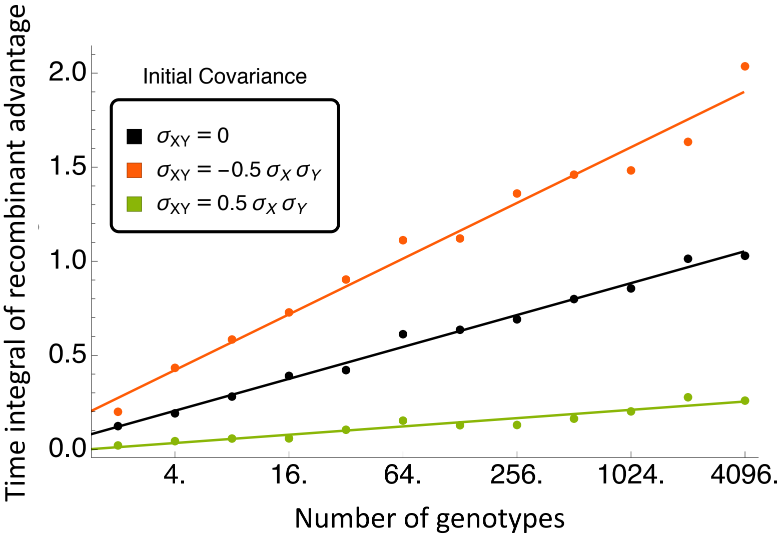


Figure S8. Time-integrated recombinant advantage increases with number of genotypes. Time-integrated recombinant advantage computed as minus one times the time-integrated covariance as predicted by Eq (S21). Values plotted were computed by Montecarlo integration, where $X$ and $Y$ were drawn from a bivariate normal distribution with $\mathbb{E}[X] = \mathbb{E}[Y] = -0.1$, and $\texttt{Var}[X] = \texttt{Var}[Y] = 0.04$, and initial covariance is indicated by color. Montecarlo integration sample size was $10,000$ for each point.

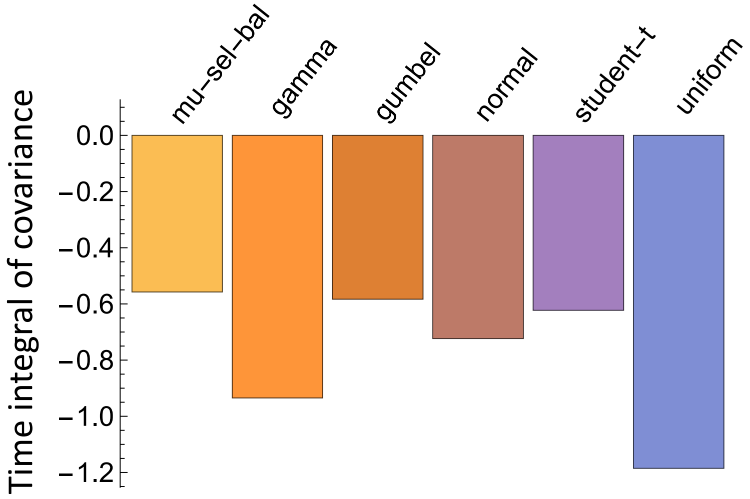


Figure S9. Time-integrated covariance as a function of distribution. Time-integrated covariance as predicted by Eq (S21). Values plotted were computed by Montecarlo integration, where $X$ and $Y$ are drawn from a bivariate distribution indicated above each bar. For all distributions, $\mathbb{E}[X] = \mathbb{E}[Y] = -0.1$, and $\texttt{Var}[X] = \texttt{Var}[Y] = 0.04$, and correlation coefficient was drawn at random from a uniform distribution over the interval $(-1, 1)$. Montecarlo integration sample size was $10,000$. With the exception of the normal distribution, all bivariate distributions were computed as bivariate Gaussian copula of two univariate distributions with CDF: $\Phi_\rho(\Phi^{-1}(F(x)), \Phi^{-1}(G(y)))$, where $\Phi_\rho$ is the bivariate standard normal CDF with correlation $\rho$, $\Phi$ is the standard normal CDF, and $F$ and $G$ are the marginal CDFs in $x$ and $y$.

#### S9. COVARIANCE DYNAMICS IN A METAPOPULATION: SIMULATIONS

**Most relevant to:** EV0 [S1] , EV1 [S3] , and EV2 [S2]

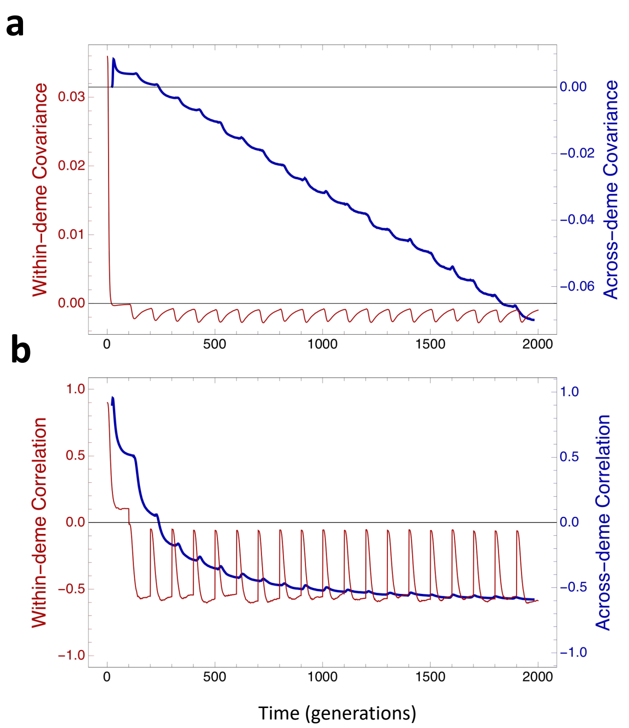


Figure S10. Covariance and correlation dynamics in a metapopulation. Simulated metapopulations of size $N = 500$ begin with all individuals being assigned unique genic fitness pairs, $(X, Y)$, drawn at random from a common bivariate normal distribution with correlation coefficient 0.9, means −0.1 and variances 0.2. Every 100 generations, uncorrelated gaussian noise was injected as follows: $X' = X + Q$ and $Y' = Y + Q$, where $Q \sim \mathcal{N}(-.1, .1)$. Plotted is **a**) mean covariance and **b**) mean correlation, where means were computed from 2000 simulations.

Detailed metapopulation simulations in which natural selection was modeled explicitly in a finite population are plotted in Fig S10. Initially, each deme of the metapopulation is assigned $n$ distinct genotypes. Natural selection then acts on this variation for 100 generations (the first bout of selection), at which time gaussian noise (mutation) was injected to initialize the next bout of selection, and the process was repeated to mimic several bouts of selection. Figure S10 reveals that within-deme (within-population) covariance becomes negative in the first bout of selection, despite starting with a correlation coefficient close to +1, as the theory in our companion paper [S2] predicts. Across-deme covariance also starts close to +1; it is reduced in the first bout of selection but does not become negative until the third bout of selection.

#### S10. ONE GENERATION OF SELECTIVE SORTING

**Most relevant to:** EV0 [S1] and EV2 [S2]

##### A. First approach

We recall that $\sigma_{XY} = \kappa_{1,1}$, $\sigma_X^2 = \kappa_{2,0}$, and $\sigma_Y^2 = \kappa_{0,2}$. We are interested in the changes in these quantities, especially in the change in $\kappa_{1,1}$ over the course of a single generation. These results are expressed most compactly in terms of cumulants, so we use the cumulant notation, recalling that $\kappa_{i,j}$ is the $(i,j)^{th}$ cumulant in $X$ and $Y$.

Change in covariance over time interval $(t, t+\delta t)$ can be written as follows:

$$\kappa_{1,1}(t+\delta t) = \kappa_{1,1}(t) + \delta t \kappa'_{1,1}(t) + \frac{1}{2}\delta t^2 \kappa''_{1,1}(t) + \mathcal{O}(\delta t^3) + ...$$

We define total change in covariance between $t = 0$ and time $t = \delta t$ to be:

$$\begin{aligned} \Delta\kappa_{1,1} &= \kappa_{1,1}(\delta t) - \kappa_{1,1}(0) \\ &= \delta t \kappa'_{1,1}(0) + \frac{1}{2}\delta t^2 \kappa''_{1,1}(0) + \mathcal{O}(\delta t^3) + ... \end{aligned} \tag{S26}$$

We will see that:

$$\kappa'_{1,1}(0) = NG_1 + D \tag{S27}$$

$$\kappa''_{1,1}(0) = NG_2 + G \tag{S28}$$

where $G$, $NG$ and $D$ refer to "gaussian", "non-gaussian" and "drift" components, respectively. We note that Eq (S27) has no gaussian component; likewise, (S28) has no drift component. Gaussian component are functions of cumulants $\kappa_{i,j}$ where $i + j \leq 2$; non-gaussian components are functions of cumulants $\kappa_{i,j}$ where $i + j > 2$.

To de-clutter the notation here, in what follows we let $\kappa_{i,j} = \kappa_{i,j}(0)$. The components come directly from Eq (S14); they are:

$$\begin{aligned} D &= -k_N \kappa_{1,1} \\ G &= -2k_N \left(4\kappa_{1,1}^2 + 3\kappa_{1,1}(\kappa_{2,0} + \kappa_{0,2}) + 2\kappa_{2,0}\kappa_{0,2}\right) \\ NG_1 &= \kappa_{1,2} + \kappa_{2,1} \\ NG_2 &= \kappa_{1,3} + 2\kappa_{2,2} + \kappa_{1,3} \end{aligned}$$

where $k_N = \frac{N-1}{N^2}$. Letting $\delta t = 1$, i.e., over the course of a single generation, we have:

$$\Delta\kappa_{1,1} = D + NG_1 + \frac{1}{2}G + \frac{1}{2}NG_2 \ ,$$

which very accurately predicts the change in covariance when compared to stochastic simulations.

For the special case where $X$ and $Y$ are initially independent, normal random variables, and the population is of infinite size, we have: $D = 0$, $NG_1 = 0$, $NG_2 = 0$, leaving:

$$\Delta\kappa_{1,1} \ = \ \frac{1}{2}G \ = \ -k_N \left(4\kappa_{1,1}^2 + 3\kappa_{1,1}(\kappa_{2,0} + \kappa_{0,2}) + 2\kappa_{2,0}\kappa_{0,2}\right) \ = \ -2k_N\kappa_{2,0}\kappa_{0,2} \ \leq \ 0.$$

Or, more generally, if covariance is initially non-negative, the action of natural selection will be to reduce the covariance ($\Delta\kappa_{1,1} \leq 0$).

We note the explicit dependence of $D$ and $G$ on population size $N$, which suggests these components can be significant in small populations but become negligible in large populations. In large populations, it is the non-gaussian components that determine the change in covariance. We also note that $D$ is a linear function in $\kappa_{1,1}$ with slope $-k_N$ and passing through the origin.

Gaussian component, $G$, is a concave parabolic function in $\kappa_{1,1}$ with maximum value:

$$\hat{G} \;=\; -k_N \left(4\kappa_{2,0}\kappa_{0,2} - \frac{9}{8}(\kappa_{2,0} + \kappa_{0,2})\right)$$

occurring at:

$$\hat{\kappa}_{1,1} = -\frac{3}{8}(\kappa_{2,0} + \kappa_{0,2})$$

If we suppose the variances in $X$ and $Y$ are equal (i.e., $\sigma_X^2 = \kappa_{2,0} = \kappa_{0,2} = \sigma_Y^2$), the foregoing maybe rewritten as maximum: $\hat{G} = \frac{1}{2}k_N\sigma_X^4$ occurring at: $\hat{\kappa}_{1,1} = \hat{\sigma}_{XY} = -\frac{3}{4}\sigma_X^2$ In this case of equal variances, the maximum should thus occur 3/4 of the way between 0 and the minimum value for covariance (in this case, the minimum value is $-\sigma_X\sigma_Y = -\sigma_X^2$). This can be seen in our simulations and in the schematic plot in Fig S11 by the fact that the G+D curve rises slightly above the D-only curve toward the negative end of the $\kappa_{1,1}$ domain and is otherwise concave, becoming increasingly smaller than D as $\kappa_{1,1}$ increases.

Of special interest is the fact that that the intercept of $G$ is unconditionally negative; i.e., when $\kappa_{1,1} = 0$, we have:

$$G(0) = -4k_N\kappa_{2,0}\kappa_{0,2} = -4k_N\sigma_X^2\sigma_Y^2$$

Add this to the fact that the intercept of $D$ is zero, and we have that when $\kappa_{1,1} = 0$:

$$G(0) + D(0) = -4k_N\sigma_X^2\sigma_Y^2 \leq 0 \tag{S29}$$

Non-gaussian component $NG_1 = \kappa_{1,2} + \kappa_{2,1}$ may be rewritten as:

$$NG_1 = \frac{1}{3}\left(\kappa_3(X+Y) - \kappa_3(X) - \kappa_3(Y)\right)$$

i.e., it is the discrepancy between the skewness of the sum and the sum of the skewnesses of $X$ and $Y$.

*Summary of results in more standard notation when the initial fitness distribution is gaussian*

The result can be partitioned into two components:

1. A "drift" component which is essentially how the covariance changes simply as a result of sampling with replacement (no selection). This is equivalent to Gibbs sampling with the Gibbs parameter equal to zero, $\beta = 0$. Letting $\Delta^d$ denote "change in $\langle$quantity$\rangle$ due to drift", we have:

$$\begin{aligned} \Delta^d\sigma_{XY} &= -k_n\sigma_{XY} \\ \Delta^d\sigma_X^2 &= -k_n\sigma_X^2 \\ \Delta^d\sigma_Y^2 &= -k_n\sigma_Y^2 \end{aligned} \tag{S30}$$

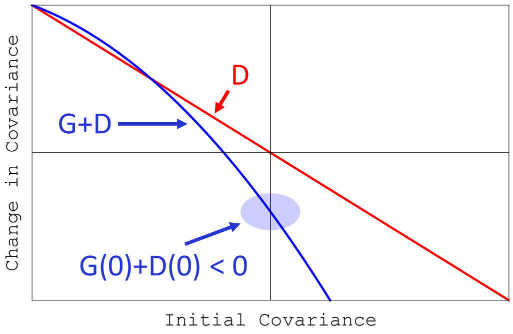


Figure S11. Schematic plot of change in covariance due to drift ($D$, in red) and drift plus the gaussian component of fitness ($G+D$, in blue) as a function of initial covariance. Three key features are: 1) Drift alone (red) is linear and symmetric passing through the origin and with an average value of zero; put differently, drift by itself has no net effect on the covariance. 2) Drift plus the gaussian component of fitness is unconditionally negative for independent $X$ and $Y$ (i.e., for initial covariance of zero); because the drift component is zero in this case, it is the gaussian component of fitness by itself that creates negative covariance. 3) On average, the gaussian component of fitness is negative; put differently, there is a net tendency to decrease covariance.

where $k_n = \frac{n-1}{n^2}$, and $n$ is population size.

2. A "selection" component, which is the change in covariance due to selection in one generation ($\beta = 1$). Letting $\Delta^s$ denote "change in $\langle$quantity$\rangle$ due to selection", we have:

$$\begin{aligned} \Delta^s \sigma_{XY} &= -k_n \left(4\sigma_{XY}^2 + 3\sigma_{XY}(\sigma_X^2 + \sigma_Y^2) + 2\sigma_X^2\sigma_Y^2\right) \\ \Delta^s \sigma_X^2 &= -k_n \left(2\sigma_{XY}^2 + \sigma_X^2(6\sigma_{XY} + 3\sigma_X^2 + \sigma_Y^2)\right) \\ \Delta^s \sigma_Y^2 &= -k_n \left(2\sigma_{XY}^2 + \sigma_Y^2(6\sigma_{XY} + 3\sigma_Y^2 + \sigma_X^2)\right) \ . \end{aligned} \tag{S31}$$

The total change in variance/covariance is the sum of these two components.

*Key observations*

1. **Independence**. A special case of interest is when $X$ and $Y$ are initially independent. This is a sort of "all else being equal" case, and it is not *a priori* obvious that covariance should decrease in this case; as will be seen, it is also an indicator of the *net* tendency. Under this condition, i.e., when covariance is initially zero, the change in covariance is:

   - Zero under drift alone:

   $$\Delta^d \sigma_{XY} = -k_n \sigma_{XY} = 0$$

   - Unconditionally negative (well, non-positive) under selection:

   $$\Delta^s \sigma_{XY} = -k_n \left(4\sigma_{XY}^2 + 3\sigma_{XY}(\sigma_X^2 + \sigma_Y^2) + 2\sigma_X^2\sigma_Y^2\right) = -2k_n\sigma_X^2\sigma_Y^2 \ \leq \ 0$$

2. **Average effects**. It is of course of interest to know the *net* or *average* effect of drift and selection on the covariance. We will let $\bar{\Delta}$ denote "average change in $\langle$quantity$\rangle$". We find the average effect to be:

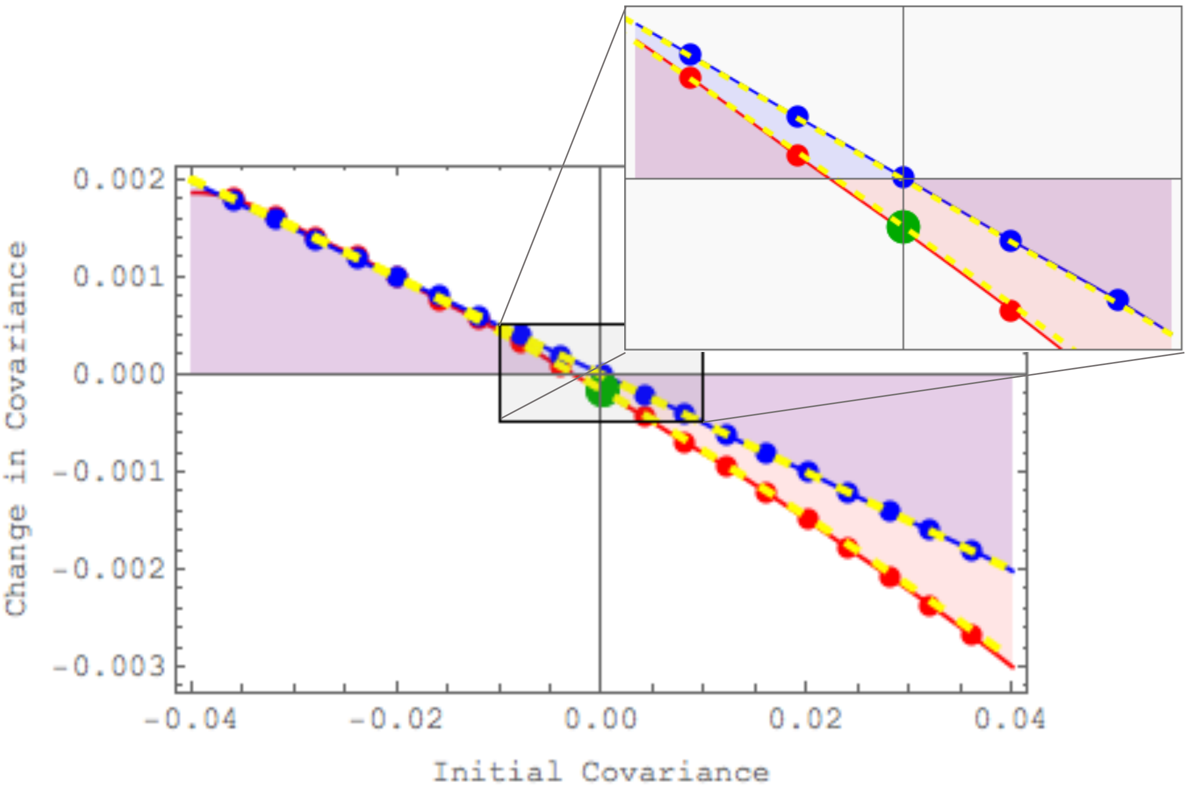


Figure S12. Change in covariance as a function of initial covariance, under: 1) drift alone ($\Delta^d\sigma_{XY}$), and 2) drift plus selection ($\Delta^d\sigma_{XY} + \Delta^s\sigma_{XY}$), within a single population. This figure plots the change in covariance due to one round of W-F (Gibbs) sampling (one generation). 1) Drift alone is in blue ($\Delta^d\sigma_{XY}$): Blue dots give the mean of one million simulations; blue curve is an interpolating function through those points. 2) Drift plus selection is in red ($\Delta^d\sigma_{XY} + \Delta^s\sigma_{XY}$): Red dots give the mean of one million simulations; red curve is an interpolating function through those points. Yellow dashed curves give theoretical predictions – Eq (S30) and the sum of Eqs (S30) and (S31) – for both curves (how about that agreement! ;-)) **Punchline:** drift alone does not give you a net downward pull on covariance; this is only achieved when selection is added. This fact is seen clearly in the surprisingly simple expressions that give the theoretical predictions: drift alone is linear and symmetric in $\sigma_{XY}$, while drift+selection is quadratic and not symmetric.

- Zero under drift alone:

$$\bar{\Delta}^d\sigma_{XY} = -\frac{k_n}{2\sigma_X\sigma_Y}\int_{-\sigma_X\sigma_Y}^{+\sigma_X\sigma_Y}\sigma_{XY}d\sigma_{XY} = 0$$

- Unconditionally negative (well, again non-positive) under selection:

$$\bar{\Delta}^s\sigma_{XY} = -\frac{k_n}{2\sigma_X\sigma_Y}\int_{-\sigma_X\sigma_Y}^{+\sigma_X\sigma_Y}\left(4\sigma_{XY}^2 + 3\sigma_{XY}(\sigma_X^2+\sigma_Y^2) + 2\sigma_X^2\sigma_Y^2\right)d\sigma_{XY} = -\frac{5}{3}k_n\sigma_X\sigma_Y \;\leq\; 0$$

#### B. Second approach

Let $X_1$, $Y_1$, $X_2$, $Y_2$ be four i.i.d. random variables with standard normal distribution. Denote:

$$Z_1 = X_1 + Y_1\ , \quad Z_2 = X_2 + Y_2;$$

$$P_1 = \frac{e^{tZ_1}}{e^{tZ_1} + e^{tZ_2}} \ , P_2 = \frac{e^{tZ_2}}{e^{tZ_1} + e^{tZ_2}} = 1 - P_1 \ ,$$

which are the frequencies of genotypes 1 and 2, respectively, at time $t$.

Let $U$ be a random variable, independent of $(X_1, Y_1, X_2, Y_2)$, uniformly distributed over $(0, 1)$. Define the random index $I \in \{1, 2\}$ as:

$$I = 1 + \mathbb{1}_{U > P_1} \ .$$

Finally, let $(X, Y) = (X_I, Y_I)$. Observe that for $t = 0$, $(X, Y)$ is distributed as $(X_1, Y_1)$ or $(X_2, Y_2)$ and therefore $X$ and $Y$ are independent. However, $\text{cov}(X, Y)$ increases for $t < 0$ and decreases for $t > 0$.

The first $\sigma$-algebra to be conditioned upon is the one generated by $(X_1, Y_1, X_2, Y_2)$ which will be denoted $\mathcal{F}_4$ as follows:

$$\begin{aligned}
\mathbb{E}[XY|\mathcal{F}_4] &= X_1 Y_1 P_1 + X_2 Y_2 P_2 \\
\mathbb{E}[X|\mathcal{F}_4] &= X_1 P_1 + X_2 P_2 \\
\mathbb{E}[Y|\mathcal{F}_4] &= Y_1 P_1 + Y_2 P_2
\end{aligned}$$

The next $\sigma$-algebra to be conditioned upon is the one generated by $(Z_1, Z_2)$ which will be denoted $\mathcal{F}_2$ as follows:

$$\begin{aligned}
\mathbb{E}[XY|\mathcal{F}_2] &= \mathbb{E}[X_1 Y_1 P_1 + X_2 Y_2 P_2 | \mathcal{F}_2] \\
&= P_1 \mathbb{E}[X_1 Y_1 | \mathcal{F}_2] + P_2 \mathbb{E}[X_2 Y_2 | \mathcal{F}_2] \ ,
\end{aligned}$$

because both $P_1$ and $P_2$ are $\mathcal{F}_2$-measurable. Now the triple $(X_1, Y_1, Z_1)$ is a Gaussian vector with null expectation, and covariance matrix:

$$\begin{pmatrix} 1 & 0 & 1 \\ 0 & 1 & 1 \\ 1 & 1 & 2 \end{pmatrix} \ .$$

Therefore the conditional distribution of $(X_1, Y_1)$ given $Z_1 = z_1$ is Gaussian with mean $(z_1, z_1)/2$ and covariance matrix:

$$\frac{1}{2} \begin{pmatrix} 1 & -1 \\ -1 & 1 \end{pmatrix}$$

Hence:

$$\begin{aligned}
\mathbb{E}[X_1 Y_1 | \mathcal{F}_2] &= \mathbb{E}[X_1 Y_1 | Z_1] \\
&= -\frac{1}{2} + \frac{1}{4} Z_1 \ .
\end{aligned}$$

Therefore:

$$\begin{aligned}
\mathbb{E}[XY|\mathcal{F}_2] &= P_1 \mathbb{E}[X_1 Y_1 | \mathcal{F}_2] + P_2 \mathbb{E}[X_2 Y_2 | \mathcal{F}_2] \\
&= P_1 \left( -\frac{1}{2} + \frac{1}{4} Z_1 \right) + P_2 \left( -\frac{1}{2} + \frac{1}{4} Z_2 \right) \\
&= -\frac{1}{2} + \frac{1}{4} \left( P_1 Z_1^2 + P_2 Z_2^2 \right) \ .
\end{aligned}$$

Now:

$$\begin{aligned}\mathbb{E}[X|\mathcal{F}_2] &= \mathbb{E}[X_1P_1 + X_2P_2|\mathcal{F}_2] \\ &= P_1\mathbb{E}[X_1|\mathcal{F}_2] + P_2\mathbb{E}[X_2|\mathcal{F}_2] \\ &= \frac{1}{2}\left(P_1Z_1 + P_2Z_2\right) .\end{aligned}$$

The same expression holds for $\mathbb{E}[Y|\mathcal{F}_2]$.

Joining together the expressions derived so far, we have:

$$\mathrm{cov}(X,Y) = -\frac{1}{2} + \frac{1}{4}\left(\mathbb{E}[P_1Z_1^2 + P_2Z_2^2] - \mathbb{E}[P_1Z_1 + P_2Z_2]^2\right) .$$

But:

$$P_1Z_1^2 + P_2Z_2^2 - (P_1Z_1 + P_2Z_2)^2 = P_1P_2(Z_1 - Z_2)^2 .$$

From there, one gets:

$$\mathrm{cov}(X,Y) = -\frac{1}{2} + \frac{1}{4}\left(\mathbb{E}\left[P_1P_2(Z_1 - Z_2)^2\right] + \mathrm{var}(P_1Z_1 + P_2Z_2)\right).$$

Remind that

$$P_1 = P_1(t) = \frac{e^{tZ_1}}{e^{tZ_1} + e^{tZ_2}} \text{ and } P_2 = P_2(t) = 1 - P_1(t).$$

The idea of the following is to study the function $f : t \in \mathbb{R} \mapsto \mathrm{cov}(X,Y)$ and its derivative. We expect that $f$ is increasing for $t < 0$ and then decreasing for $t > 0$ namely $\mathrm{Sign}(f'(t)) = -\mathrm{Sign}(t)$. We have

$$f'(t) = \frac{1}{4}\mathbb{E}\left[(P_1'P_2 + P_1P_2')(Z_1 - Z_2)^2\right] + \frac{1}{2}\mathrm{cov}(P_1'Z_1 + P_2'Z_2, P_1Z_1 + P_2Z_2), \tag{S32}$$

where the second term in the RHS can be seen as the derivative of the symmetric bilinear form $\mathrm{var}(P_1Z_1 + P_2Z_2) = \mathrm{cov}(P_1Z_1 + P_2Z_2, P_1Z_1 + P_2Z_2)$.

**First term in** (S32). The first term in Equation (S32) is of the sign of $-t$. Indeed, remark that $P_1'P_2 + P_1P_2' = P_1'(1 - P_1) + P_1(-P_1') = P_1'(1 - 2P_1)$,

$$(1 - 2P_1(t)) = \frac{e^{tZ_2} - e^{tZ_1}}{e^{tZ_1} + e^{tZ_2}}$$

and

$$P_1'(t) = \frac{Z_1e^{tZ_1}(e^{tZ_1} + e^{tZ_2}) - e^{tZ_1}(Z_1e^{tZ_1} + Z_2e^{tZ_2})}{(e^{tZ_1} + e^{tZ_2})^2} = \frac{(Z_1 - Z_2)e^{t(Z_1+Z_2)}}{(e^{tZ_1} + e^{tZ_2})^2}.$$

We deduce that $\mathrm{Sign}(1 - 2P_1(t)) = \mathrm{Sign}(t(Z_2 - Z_1))$ and $\mathrm{Sign}(P_1'(t)) = \mathrm{Sign}(Z_1 - Z_2)$ hence

$$\mathrm{Sign}\left(P_1'(t)(1 - 2P_1(t))\right) = -\mathrm{Sign}(t).$$

As a particular case, this term is null when $t = 0$.

**Second term in** (S32). As a first step I tried to evaluate it when $t = 0$. Notice that $P_1(0) = 1/2$ and $P_1'(0) =$

$(Z_1 - Z_2)/4$ so that it rewrites as

$$\begin{aligned}\operatorname{cov}(P_1' Z_1 + P_2' Z_2, P_1 Z_1 + P_2 Z_2) &= \operatorname{cov}\left(\frac{(Z_1 - Z_2)^2}{4}, \frac{Z_1 + Z_2}{2}\right) \\ &= \mathbb{E}\left[\frac{(Z_1 - Z_2)(Z_1^2 - Z_2^2)}{8}\right] - \mathbb{E}\left[\frac{(Z_1 - Z_2)^2}{4}\right]\mathbb{E}\left[\frac{Z_1 + Z_2}{2}\right] \\ &= \frac{1}{8}\mathbb{E}\left[Z_1^3 + Z_2^3\right] - 0.\end{aligned}$$

where we assumed in the last line that the variables are centred.

**Summary.** Hence $f'(0) = \mathbb{E}\left[Z_1^3\right]/4$ which is non null in general. It suggests that for $|t|$ small enough we can recover some positive covariance. If the third moment/skewness is negative then it happens for negative $t$'s and conversely when the skewness is positive.

When $Z$ is a centered normal distribution, $\mathbb{E}[Z^3] = 0$ so that $f'(0) = 0$. However, we find that

$$f''(0) \;=\; -\frac{1}{8}\mathbb{E}[(X_1 - X_2)(Y_1 - Y_2)(Z_1 - Z_2)^2]$$

When $(X_1, Y_1, X_2, Y_2)$ are i.i.d. standard normal,

$$f''(0) \;=\; -1\ ,$$

from which, together with $f(0) = f'(0) = 0$, we infer that natural selection causes covariance to become negative.

#### S11. WHEN RESIDENT POPULATION ALREADY HAS SOME LEVEL OF RECOMBINATION

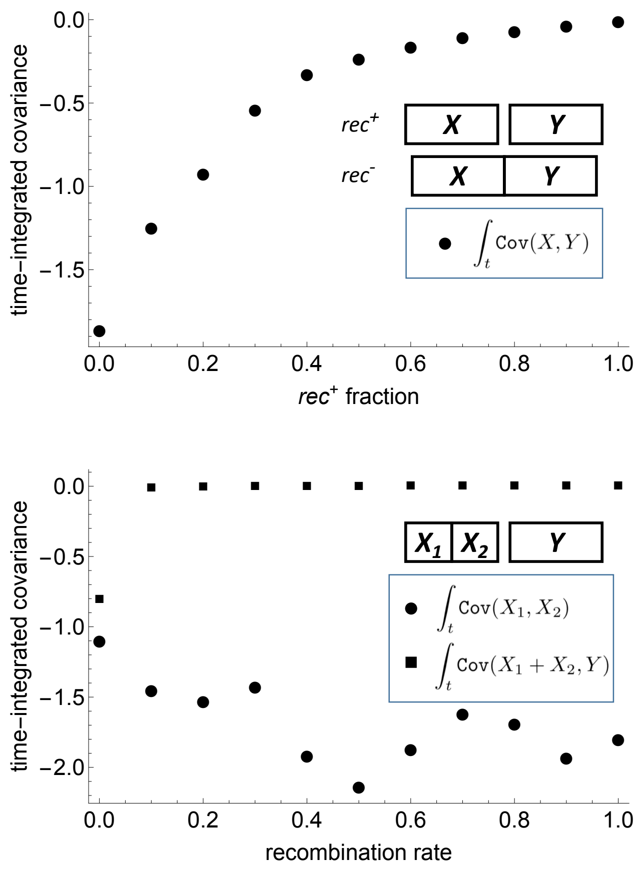


Figure S13. Selective value of further recombination when some level of recombination is already present. Top panel: time-integrated covariance when a specified fraction of the resident population (horizontal axis) is already $rec^+$. Bottom panel: time-integrated covariance when two loci, $X1$ and $X2$, are linked but recombination already occurs between the $X1/X2$ block and the $Y$ locus. Negative covariance builds up within the $X1/X2$ block. The evolution of recombination withing the $X1/X2$ block is thus promoted. $X$ and $Y$ were drawn from a bivariate normal distribution with $\mathbb{E}[X] = \mathbb{E}[Y] = -0.1$, and $\mathtt{Var}[X] = \mathtt{Var}[Y] = 0.04$, and initial covariance is zero. Each point is the average of 2000 simulations.

#### S12. RELATION TO TWO GENERAL CONDITIONING EFFECTS OF PARTICULAR RELEVANCE

Our study identifies phenomena that are inherent consequences of natural selection and give rise to negative fitness associations across loci. This pervasive phenomenon is an example of counter-intuitive effects caused by probabilistic conditioning quite generally. Two examples of special relevance here are "Berkson's paradox" [S7, S8] and the "Bulmer effect" [S9–S14].

##### A. Berkson's paradox

Berkson's paradox arises when a selective observational procedure produces spurious negative correlations. In the original context, among those admitted to hospital, it is disproportionately more likely for a patient to have, say, diabetes or, say, hypertension than to have both. Observed negative correlation between diabetes and hypertension can therefore be an artefact created by hospital admission and not representative of the general population. The selective agent in this case is hospital admission. The negative correlation may also be understood by considering that people with neither diabetes nor hypertension are less likely to be admitted to the hospital than people chosen at random from the general population.

To make the above example numerical and concrete, let us suppose that $A$ denotes the event that a person is diabetic and that $B$ denotes the event that a person is hypertensive. Let us further suppose that:

- half of the general population is diabetic ($\mathbb{P}(A) = 1/2$)
- half of the general population is hypertensive ($\mathbb{P}(B) = 1/2$)
- incidences of diabetes and hypertension are (unrealistically) independent ($\mathbb{P}(AB) = \mathbb{P}(A)\mathbb{P}(B) = 1/4$)
- all people who have one or the other condition are admitted to hospital

In a general population of 100 therefore we have on average the incidence table shown in Fig S14, illustrating how the sample space is reduced by conditioning:

| | $A$ | $\bar{A}$ |
|---|---|---|
| $B$ | 25 | 25 |
| $\bar{B}$ | 25 | 25 |

Figure S14. Numerical example of Berkson's paradox. A population of 100 is equally distributed into the four categories. $A$ and $!A$ denote diabetic and non-diabetic, respectively. $B$ and $!B$ denote hypertensive and non-hypertensive, respectively. Categories highlighted in yellow are those admitted to hospital. Put differently, yellow highlights the set $A \cup B$.

*Sample space reduction:*

- General population: 100
- Those admitted to hospital: 75

This example reveals how negative associations arise as a consequence of the reduction in sample size:

- In general population:

$$\texttt{Cov}(A,B) \ = \ \mathbb{P}(AB) - \mathbb{P}(A)\mathbb{P}(B) \ = \ \frac{25}{100} - \frac{50}{100} \times \frac{50}{100} \ = \ \frac{1}{4} - \frac{1}{2} \times \frac{1}{2} \ = \ 0 \ ,$$

as expected from our assumption of independence.

- Of those admitted to hospital:

$$\texttt{Cov}^*(A,B) \ = \ \mathbb{P}(AB|A \cup B) - \mathbb{P}(A|A \cup B)\mathbb{P}(B|A \cup B) \ = \ \frac{25}{75} - \frac{50}{75} \times \frac{50}{75} \ = \ \frac{1}{3} - \frac{2}{3} \times \frac{2}{3} \ = \ -\frac{1}{9} \ < \ 0 \ .$$

In general, we have:

→ $\mathbb{P}(AB|A \cup B) = \mathbb{P}(AB)/(1 - \mathbb{P}(\bar{A}\bar{B}))$

→ $\mathbb{P}(A|A \cup B) = \mathbb{P}(A)/(1 - \mathbb{P}(\bar{A}\bar{B}))$

→ $\mathbb{P}(B|A \cup B) = \mathbb{P}(B)/(1 - \mathbb{P}(\bar{A}\bar{B}))$

giving:

$$\texttt{Cov}^*(A,B) \ = \ \mathbb{P}(AB)/(1 - \mathbb{P}(\bar{A}\bar{B})) - \mathbb{P}(A)\mathbb{P}(A)/(1 - \mathbb{P}(\bar{A}\bar{B}))^2$$

When incidence of diabetes and hypertension are independent in the general population (as we have rather unrealistically assumed), then:

$$\texttt{Cov}^*(A,B) \ = \ \mathbb{P}(A)\mathbb{P}(B)\left(1/(1 - \mathbb{P}(\bar{A}\bar{B})) - 1/(1 - \mathbb{P}(\bar{A}\bar{B}))^2\right)$$

Noting that $\mathbb{P}(A)\mathbb{P}(B) \geq 0$, $\mathbb{P}(\bar{A}\bar{B}) \geq 0$ and:

$$1/(1 - \mathbb{P}(\bar{A}\bar{B})) - 1/(1 - \mathbb{P}(\bar{A}\bar{B}))^2 \ = \ -\mathbb{P}(\bar{A}\bar{B})/(1 - \mathbb{P}(\bar{A}\bar{B}))^2 \ \leq \ 0$$

yields:

$$\texttt{Cov}^*(A,B) \ \leq \ 0$$

##### B. The Bulmer effect

Negative correlations can also arise across genetically-encoded trait values as a consequence not of selective observation (as in Berkson's paradox) but of selective sorting. And as in Berkson's paradox, this results in part because genotypes in which both loci have low genic fitness are purged by selection, whether selection be natural or artificial; in this case, however, the resulting bias is not observational but actual, as these low-fitness genotypes are selectively removed from the population. In this genetic context, where selection is imposed artificially through breeding, this biasing effect has been contemplated and modeled; it is known as the "Bulmer effect" [S9–S14]. In its original formulation, a desired (artificially selected) trait is determined by several genetic components; one generation of breeding for the desired trait will result in negative associations among the trait's components. The focus of Bulmer's original work was not the negative associations created across loci but the reduction in trait variation that resulted.

We now present a simple model that will: 1) help to gain an intuitive understanding of the Bulmer effect, and 2) illustrate how the Bulmer effect is a sort of genetic Berkson's paradox (extending the Berkson's paradox model).

Our model is a two-locus, two-allele model with four genotypes $ab$, $Ab$, $aB$, and $AB$. Lower-case letters denote wildtype alleles with multiplicative fitness contributions equal to one. Upper-case letters denote alleles with fitnesses increased by factors $1 + s_A$ and $1 + s_B$. To simplify presentation, we will assume that these factors are the same such that $s_A = s_B = s$. In the context of the Bulmer effect, fitness is due to artificial selection, but this fact is immaterial to the present model (i.e., it could just as well be natural selection).

Fitnesses of the four possible genotypes are thus given by the table in Fig S15.

| | A | a |
|---|---|---|
| B | $w_{AB}= (1+s)^2$ | $w_{aB}= (1+s)$ |
| b | $w_{Ab}= (1+s)$ | $w_{ab}= 1$ |

Figure S15. Fitnesses of four possible genotypes in simple model of the Bulmer effect.

Frequencies of the four possible genotypes are given by the table in Fig S16.

| | A | a |
|---|---|---|
| B | f(AB) | f(aB) |
| b | f(Ab) | f(ab) |

Figure S16. Fitnesses of four possible genotypes in simple model of the Bulmer effect. Marginal frequencies of the higher fitness alleles are: $f(A) = f(AB) + f(Ab)$ and $f(B) = f(AB) + f(aB)$.

*Before selection (general population)*

Let random variables $X$ and $Y$ denote genic fitnesses at the first and second loci. Then we have:

$$\begin{aligned}\mathbb{E}[XY] &= f(AB)(1+s)^2 + f(Ab)(1+s) + f(aB)(1+s) + f(ab)\\ \mathbb{E}[X] &= f(A)(1+s) + f(a)\\ \mathbb{E}[Y] &= f(B)(1+s) + f(b)\end{aligned}$$

where $f(A) = f(AB) + f(Ab)$, $f(B) = f(AB) + f(aB)$, $f(a) = f(aB) + f(ab)$, and $f(b) = f(Ab) + f(ab)$. From here, we have genic fitness covariance for the general population:

$$\begin{aligned}\texttt{Cov}(X,Y) &= \mathbb{E}[XY] - \mathbb{E}[X]\mathbb{E}[Y]\\ &= [f(AB) - f(A)f(B)]s^2\end{aligned}$$

When events $A$ and $B$ are independent, we have $f(AB) = f(A)f(B)$, and:

$$\texttt{Cov}(X,Y) = 0 \ ,$$

as expected.

*After selection*

Here, we assume a very simple form of selection: we assume that only individuals with fitness greater than one survive (through "culling" or truncation selection). After selection, therefore, our sample space is reduced from $\{AB, Ab, aB, ab\}$ to $\{AB, Ab, aB\}$. As a result, we now have:

$$\begin{aligned}\mathbb{E}[XY] &= [f(AB)(1+s)^2 + f(Ab)(1+s) + f(aB)(1+s)]/(1-f(ab)) \\ \mathbb{E}[X] &= [f(A)(1+s) + f(a)]/(1-f(ab)) \\ \mathbb{E}[Y] &= [f(B)(1+s) + f(b)]/(1-f(ab))\end{aligned}$$

where the marginal frequencies are now: $f(A) = f(AB) + f(Ab)$, $f(B) = f(AB) + f(aB)$, $f(a) = f(aB)$, and $f(b) = f(Ab)$. From here, we have genic fitness covariance for the "culled" or "truncated" population (i.e., after selection):

$$\begin{aligned}\mathtt{Cov}^*(X,Y) &= \mathbb{E}[XY] - \mathbb{E}[X]\mathbb{E}[Y] \\ &= \frac{f(AB)(1+s)^2 + f(Ab)(1+s) + f(aB)(1+s)}{1-f(ab)} - \frac{(f(A)(1+s)+f(a))(f(B)(1+s)+f(b))}{(1-f(ab))^2}\end{aligned}$$

Before selection, we assume independence between the two loci such that $f(ij) = f(i)f(j)$, which gave pre-selection covariance $\mathtt{Cov}(X,Y) = 0$ as expected. Starting with independence before selection, the post-selection covariance between genic fitnesses reduces to:

$$\mathtt{Cov}^*(X,Y) = \left(\Theta^{-1} - \Theta^{-2}\right) f(A)f(B)s^2$$

where $\Theta = f(A) + f(B) - f(A)f(B)$. By virtue of the facts that $0 \leq f(A) \leq 1$ and $0 \leq f(B) \leq 1$, we have that $f(A) \geq f(A)f(B)$ and $f(B) \geq f(A)f(B)$, giving $\Theta \leq 1$ and consequently $\Theta^{-1} - \Theta^{-2} \leq 0$; hence, the post-selection covariance satisfies:

$$\mathtt{Cov}^*(X,Y) \leq 0$$

If we let $Z = X + Y$, then Bulmer was more in the fact that pre-selection variance is $\mathtt{Var}(Z) = \mathtt{Var}(X) + \mathtt{Var}(Y)$, and post-selection variance is $\mathtt{Var}^*(Z) = \mathtt{Var}(X) + \mathtt{Var}(Y) + 2\mathtt{Cov}^*(X,Y)$, which gives the relation:

$$\mathtt{Var}^*(Z) \leq \mathtt{Var}(Z)$$

by virtue of the fact that $\mathtt{Cov}^*(X,Y) \leq 0$. Bulmer's focus was not the negative covariance between trait components $X$ and $Y$ but the reduction in variance of the trait $Z$.

On the surface, the Bulmer effect appears similar to the phenomena we describe. On closer examination, however, it differs in several critical ways: 1) Bulmer was concerned primarily with the immediate effect of selection on trait variance, whereas our analyses focus on the long-term effect of selection on the selective value of recombination, 2) the Bulmer effect requires that the components of the selected trait must each be highly multigenic such that the central limit theorem applies (i.e., fitness components upon which natural selection acts are multivariate normal), whereas our model has no such requirement, 3) the Bulmer effect requires that the pre-selection state of the components must be independent (i.e., zero covariance in the multivariate normal), whereas our general model does not make this assumption, 4) the negative association arising from the Bulmer effect quickly disappears in subsequent generations because free recombination is assumed, whereas our model examines the effects of a wide range of linkage – from fleeting to sustained – on the evolution of recombination, 5) Bulmer's model does not specify the number of components contributing to the selected trait but only gives the total reduction in variance incurred by the negative associations; as such it gives no information about the selective value of individual recombinants or recombination modifiers. In general, our approach is well-suited to studying the evolution of sex and recombination under very general conditions, whereas the Bulmer effect is more about an inefficiency in artificial selection in freely recombining populations under fairly restrictive conditions. Finally, in developments of the preceding section (that are somewhat peripheral to the main focus of our work), we have proven that when the components of fitness are completely linked and are multivariate normal with zero covariance, the immediate effect of selection is to create negative covariance among the components, in expectation – a sort of Bulmer effect for the case of complete linkage instead of free recombination.

#### S13. SOME OBSERVATIONS, DIRECTIONS AND QUESTIONS FOR FURTHER STUDY

**Most relevant to:** EV0 [S1] and EV1 [S3]

##### A. Negative association

The original reference on negative association is Joag-Dev and Proschan [S15]. The theory is briefly sketched in some textbooks, like in section 3.4 p. 148 of [S16], or section 3.3 p. 43 of [S17]. Useful reviews have been given by Pemantle [S18], and Borcea et al. [S19]. Sufficient conditions for negative dependence in random vectors have been investigated in [S20, S21]. We shall use the conventions of the above references, and call "increasing" a function from $\mathbb{R}^k$ into $\mathbb{R}$ which is nondecreasing in each coordinate. Here is the definition of negative association, such as given for instance in [S17, p. 46].

Definition S13.1. *A random vector $Y = (Y_1, \ldots, Y_m)$ is called* Negatively Associated *(NA) if for every pair $J_1, J_2$ of disjoint subsets of $\{1, \ldots, m\}$, and for every pair $f_1, f_2$ of real-valued increasing functions:*

$$\mathrm{cov}(f_1(Y_{j_1}, j_1 \in J_1), f_2(Y_{j_2}, j_2 \in J_2)) \leqslant 0 \ .$$

Intuitively, a vector is NA if when some coordinates are raised, the others must be lowered on average.

Here are two tentative definitions for fitness and recombination.

Definition S13.2. *Let $Y = (Y_1, \ldots, Y_m)$ be a $m$-dimensional random vector. The* fitness *of $Y$ is the exponential of the sum of coordinates.*

$$\mathrm{fit}(Y) = \exp\left(\sum_{j=1}^{m} Y_j\right) \ .$$

Definition S13.3. *Let $Y = (Y_1, \ldots, Y_m)$ be a $m$-dimensional random vector, and let $Y^*$ be an independent copy of $Y$. Let $J \subset \{1, \ldots, m\}$ be a set of indices. The* recombination of $Y$ over $J$ *is the random vector $Y^J = (Y_1^J, \ldots, Y_m^J)$ constructed as follows.*

$$Y_j^J = \begin{cases} Y_j & \text{if } j \in J \\ Y_j^* & \text{else.} \end{cases}$$

The interest of negative association lies in the simple observation that if a random vector is NA, then any recombination has a better expected fitness.

Proposition S13.1. *Let $Y = (Y_1, \dots, Y_m)$ be a $m$-dimensional random vector, and assume it is NA. Then for all $J \subset \{1, \dots, m\}$,*

$$\mathbb{E}(\text{fit}(Y)) \leqslant \mathbb{E}(\text{fit}(Y^J)) .$$

*Proof.* Let $Y^*$ be an independent copy of $Y$, as in Definition S13.3. Since $Y$ and $Y^*$ are i.i.d.,

$$\mathbb{E}(\text{fit}(Y^J)) = \mathbb{E}\left(\exp\left(\sum_{j_1 \in J} Y_{j_1}\right)\right) \mathbb{E}\left(\exp\left(\sum_{j_2 \notin J} Y_{j_2}\right)\right) .$$

Thus:

$$\text{fit}(Y) - \text{fit}(Y^J) = \text{cov}\left(\exp\left(\sum_{j_1 \in J} Y_{j_1}\right), \exp\left(\sum_{j_2 \notin J} Y_{j_2}\right)\right) .$$

If $Y$ is NA, the above covariance is negative or null: this is a particular case of Definition S13.1, taking $J_1 = J$, $J_2 = \{1, \dots, m\} \setminus J$,

$$f_1(Y_{j_1}, j_1 \in J_1) = \exp\left(\sum_{j_1 \in J} Y_{j_1}\right) \quad \text{and} \quad f_2(Y_{j_2}, j_2 \in J_2) = \exp\left(\sum_{j_2 \notin J} Y_{j_2}\right) .$$

□

In view of Proposition S13.1, the question is: which conditions imply that a vector of gene contributions is NA? It is reasonable to expect that negative association should arise from conditioning. Indeed consider independent coordinates $X_1, \dots, X_m$, an increasing function $\phi$ from $\mathbb{R}^m$ into $\mathbb{R}$, and some domain $\Delta \subset \mathbb{R}$. As remarked by Hu and Yang [S21], the conditional distribution of $(X_1, \dots, X_m)$ given $\phi(X_1, \dots, X_m) \in \Delta$ should be NA under mild conditions over $\Delta$. Somewhat surprisingly, rigorous results in that direction are scarce. Of most interest to us is the following basic fact, stated in [S17, p. 48]. The proof can be found in [S15].

Theorem S13.1. *Let $(X_1, \dots, X_m)$ be a vector of independent random variables, each with a logconcave density (that is, Polya frequency of order 2, or $PF_2$). Then the joint conditional distribution of $(X_1, \dots, X_m)$ given their sum is NA.*

It is reasonable to expect that conditioning over an event depending on the sum, like "the sum ranks $i$-th" should yield an analogous conclusion.

Conjecture S13.4. *With the notations of section S3 A, the ranking function being the sum of coordinates, assume that the coordinates of $X$ are independent with $PF_2$ densities. Then $X_{(i)}$ is NA, for all $i = 1, \dots, n$.*

In order to understand how Conjecture S13.4 relates to Theorem S13.1, we need to express covariances of functions of $X_{(i)}$ in terms of conditional distributions of $X$ given the sum of its coordinates. This will be done through the following technical Lemma on conditional covariances: see [S16, p. 149].

Lemma S13.1. *Let $(X, Y)$ be a pair of real-valued random variables on $\Omega$, and let $\mathcal{F}_1 \subseteq \mathcal{F}_2$ be two $\sigma$-fields on $\Omega$. Then:*

$$\text{cov}\,(X, Y \,|\, \mathcal{F}_1) = \mathbb{E}\left(\text{cov}\,(X, Y \,|\, \mathcal{F}_2) \,|\, \mathcal{F}_1\right) + \text{cov}\left(\mathbb{E}(X \,|\, \mathcal{F}_2)\,, \mathbb{E}(Y \,|\, \mathcal{F}_2) \,|\, \mathcal{F}_1\right) .$$

*In particular, when* $\mathcal{F}_1 = \{\emptyset, \Omega\}$*:*

$$\mathrm{cov}\,(X,Y) = \mathbb{E}\,(\mathrm{cov}\,(X,Y \,|\, \mathcal{F}_2)) + \mathrm{cov}\,(\mathbb{E}(X \,|\, \mathcal{F}_2)\,,\, \mathbb{E}(Y \,|\, \mathcal{F}_2))\ .$$

PROPOSITION S13.2. *With the notations of sections S3A and S3B, let* $f_1$ *and* $f_2$ *be two real-valued functions of* $m$ *variables. Then:*

$$\begin{aligned}\mathrm{cov}(f_1(X_{(i)}), f_2(X_{(i)})) = &\int_{\mathbb{R}} \mathrm{cov}(f_1(X_1), f_2(X_1) \,|\, \varphi_1 = x)\, \mathrm{d}H_i(x) \\ &+ \mathrm{cov}(\mathbb{E}(f_1(X_1) \,|\, \varphi_1), \mathbb{E}(f_2(X_1) \,|\, \varphi_1) \,|\, \sigma(i) = 1)\ .\end{aligned}$$

*Proof.* Recall from Lemma S3.1 that the distribution of $X_{(i)}$ is the conditional distribution of $X_1$ given $\sigma(i) = 1$.

$$\mathrm{cov}(f_1(X_{(i)}), f_2(X_{(i)})) = \mathrm{cov}(f_1(X_1), f_2(X_1) \,|\, \sigma(i) = 1)\ .$$

Define as $\mathcal{F}_1$ the $\sigma$-algebra generated by the event $\sigma(i) = 1$. Let $\mathcal{F}_2$ be generated by $\mathcal{F}_1$ and $\varphi_1$. Lemma S13.1 translates as:

$$\begin{aligned}\mathrm{cov}(f_1(X_{(i)}), f_2(X_{(i)})) = &\ \mathbb{E}(\mathrm{cov}(f_1(X_1), f_2(X_1) \,|\, \varphi_1) \,|\, \sigma(i) = 1) \\ &+ \mathrm{cov}(\mathbb{E}(f_1(X_1) \,|\, \varphi_1), \mathbb{E}(f_2(X_1) \,|\, \varphi_1) \,|\, \sigma(i) = 1)\ .\end{aligned}$$

To transform the first term in the right handside, observe that given the value of $\varphi_1$, $X_1$ is independent from the event $\sigma(i) = 1$. □

Assume now that $f_1$ and $f_2$ are increasing, and depend on disjoint subsets of coordinates. If the hypothesis of Conjecture S13.4 holds, then by Theorem S13.1, for all $x$:

$$\mathrm{cov}(f_1(X_1), f_2(X_1) \,|\, \varphi_1 = x) \leqslant 0\ .$$

However,

$$\mathrm{cov}(\mathbb{E}(f_1(X_1) \,|\, \varphi_1), \mathbb{E}(f_2(X_1) \,|\, \varphi_1) \,|\, \sigma(i) = 1) \geqslant 0\ ,$$

because both $\mathbb{E}(f_1(X_1) \,|\, \varphi_1)$ and $\mathbb{E}(f_2(X_1) \,|\, \varphi_1)$ are nondecreasing functions of $\varphi_1$. Thus Proposition S13.2 expresses $\mathrm{cov}(f_1(X_{(i)}), f_2(X_{(i)}))$ as the sum of two terms of opposite sign, and there does not seem to be any obvious reason why the first (negative) term should dominate.

#### B. Stochastic monotonicity

Due to their interest in reliability theory, a huge literature has developed around the many notions of stochastic ordering of random variables and random vectors. We shall use here only the so called "usual multivariate stochastic order", such as described in section 6.B of [S22, p. 266].

DEFINITION S13.5. *Let* $m$ *be an integer,* $X, Y$ *be two* $m$*-dimensional random vectors. The random vector* $X$ *is stochastically dominated by* $Y$*, or* $X \leqslant_{\mathrm{st}} Y$ *if*

$$\mathbb{E}(\psi(X)) \leqslant \mathbb{E}(\psi(Y))\ ,$$

*for all increasing function $\psi$ from $\mathbb{R}^m$ into $\mathbb{R}$, such that both expectations exist.*

As in the univariate case a coupling characterization is available [S22, p. 267]. With the notations of section S3 A, obviously:

$$\phi(X_{(1)}) \leqslant_{\text{st}} \cdots \leqslant_{\text{st}} \phi(X_{(n)}) \; .$$

If the ranking function $\phi$ is increasing, it is natural to expect that $X_{(1)}, \ldots, X_{(n)}$ are also stochastically ordered, and that this can be deduced from existing results without any further hypothesis. This has proved frustratingly elusive.

Conjecture S13.6. *Assume the ranking function $\phi$ is increasing. Then $X_{(i)}$ is stochastically increasing in $i$.*

The only rigorous result we have been able to come up with in this direction, uses the pdf calculated in Proposition S3.1.

Proposition S13.3. *Assume that the ranking function $\phi$ is increasing, and that the distribution of $X$ is (positively) associated. Assume moreover that the hypothesis of Proposition S3.1 holds. Then:*

$$X_{(1)} \leqslant_{\text{st}} X \leqslant_{\text{st}} X_{(n)} \; .$$

*Proof.* Theorem 6.B.8 in [S22, p. 270] states that if $X$ is associated (which holds in particular if the coordinates are independent), and the ratio of the density of $Y$ by the density of $X$ is increasing, then $X \leqslant_{\text{st}} Y$. By Proposition S3.1, the ratio of the density of $X_{(n)}$ by that of $X$ is $nH^{n-1}(\phi(x_1, \ldots, x_m))$, which is indeed increasing. Hence $X \leqslant_{\text{st}} X_{(n)}$. For the other inequality, replace $X$ by $-X$. □

The basic result on conditioning by sums, originally due to Efron [S23] is Theorem 6.B.9 [S22, p. 270]. Before stating the result, the authors remark that if $\phi$ is an increasing $m$-dimensional function, it seems reasonable to expect that the conditional distribution of a vector $X$ given $\phi(X) = \varphi$, should be stochastically increasing in $\varphi$. However, this is not always true. It is true when the coordinates of $X$ are independent and $\phi$ is the sum, with an additional technical hypothesis on the densities.

Theorem S13.2. *With the notations of section S3 A, the ranking function $\phi$ being the sum, assume that the coordinates of $X$ are independent, each with $PF_2$ density. Then the conditional distribution of $X$ given $\phi(X) = \varphi$ is stochastically monotone in $\varphi$.*

Notice the analogy of the previous statement with Theorem S13.1 on negative association. A major difference is that the usual stochastic order is closed under mixtures, whereas no such result holds for negative association. Theorem S13.3 below is statement e) of Theorem 6.B.16 in [S22, p. 273].

Theorem S13.3. *Let $X$, $Y$, and $\Theta$ be random vectors such that for all $\theta$ in the support of $\Theta$:*

$$[X \,|\, \Theta = \theta] \leqslant_{\text{st}} [Y \,|\, \Theta = \theta] \; .$$

*Then $X \leqslant_{\text{st}} Y$.*

We have failed to see precisely how to relate Theorems S13.2 and S13.3 to Conjecture S13.6. However, a similar relation is made by Khaledi and Kochar [S24] who investigate, in the context of concomitants of order statistics, the different order relation that may arise from positive dependence assumptions between the concomitants and the ranked variable. In particular, point (a) of their Theorem 3.1 [S24, p. 269] states something like "if $X$ is stochastically increasing in $\phi$ (like Theorem S13.2 says), then the concomitants are stochastically increasing in $i$ (like Conjecture S13.6)". However concomitants there are considered by Khaledi and Kochar in the bivariate case. Also, they assume that the conditional distributions have densities, which does not hold here. That their proof can be adapted to our setting remains to be checked. Concomitants are reviewed in section 6.8 of [S25, p. 144]. The multivariate generalization is treated on [S25, p. 146]. Khaledi and Kochar's article has a follow-up by Blessinger [S26]. Two more recent papers which appear to be related are [S27] and [S28].

A natural intuition behind Conjecture S13.4, is that conditioning makes covariances smaller. Something like the following statement.

Conjecture S13.7. *Let $f_1$ and $f_2$ be two increasing functions of $m$ variables. Then for all $i = 1, \dots, n$:*

$$\mathrm{cov}(f_1(X), f_2(X)) \geqslant \mathrm{cov}(f_1(X_{(i)}), f_2(X_{(i)})) \,.$$

Indeed, if the coordinates of $X$ are independent and $f_1$, $f_2$ depend on disjoint subsets of coordinates, then $\mathrm{cov}(f_1(X), f_2(X)) = 0$. Thus Conjecture S13.7 would imply Conjecture S13.4. The only rigorous result we have been able to come up with in this direction, is the following Proposition.

Proposition S13.4. *Let $f_1$ and $f_2$ be two real-valued functions of $m$ variables. Provided all expectations written below are finite,*

$$\begin{aligned} \frac{1}{n}\sum_{i=1}^{n} \mathrm{cov}(f_1(X_{(i)}), f_2(X_{(i)})) &= \mathrm{cov}(f_1(X), f_2(X)) \\ &- \frac{1}{2n^2}\sum_{i_1,i_2=1}^{n} \big(\mathbb{E}(f_1(X_{(i_1)})) - \mathbb{E}(f_1(X_{(i_2)}))\big) \\ &\qquad \times \big(\mathbb{E}(f_2(X_{(i_1)})) - \mathbb{E}(f_2(X_{(i_2)}))\big) \,. \end{aligned} \tag{S33}$$

Let $f_1$ and $f_2$ be two *increasing* functions. If Conjecture S13.6 holds, both sequences $(\mathbb{E}(f_1(X_{(i)}))_{1\leqslant i\leqslant n}$ and $(\mathbb{E}(f_2(X_{(i)}))_{1\leqslant i\leqslant n}$ are nondecreasing. This implies that the sum in the right handside of (S33) is nonnegative (Chebyshev's sum inequality). Therefore:

$$\frac{1}{n}\sum_{i=1}^{n} \mathrm{cov}(f_1(X_{(i)}), f_2(X_{(i)})) \leqslant \mathrm{cov}(f_1(X), f_2(X)) \,.$$

Unfortunately, even if this were proved, it would not imply that any particular covariance in the sum should be smaller than $\mathrm{cov}(f_1(X), f_2(X))$.

*Proof.* Consider a random variable $I$, independent from $X_1, \dots, X_n$, uniformly distributed over $\{1, \dots, n\}$. As already remarked in section S3 B, choosing at random one of the $X_{(i)}$'s is equivalent to choosing any of the $X_i$'s. Thus $X_{(I)}$ and $X$ have the same distribution. In particular:

$$\mathrm{cov}(f_1(X_{(I)}), f_2(X_{(I)})) = \mathrm{cov}(f_1(X), f_2(X)) \,.$$

Apply Lemma S13.1 by conditioning over $I$:

$$\begin{aligned} \mathrm{cov}(f_1(X_{(I)}), f_2(X_{(I)})) &= \mathbb{E}(\mathrm{cov}(f_1(X_{(I)}), f_2(X_{(I)}))\,|\,I) \\ &\quad + \mathrm{cov}(\mathbb{E}(f_1(X_{(I)}\,|\,I), \mathbb{E}(f_2(X_{(I)}\,|\,I)) \,. \end{aligned}$$

On the right-hand side of the above identity, the first expectation is the arithmetic mean:

$$\frac{1}{n}\sum_{i=1}^{n} \mathrm{cov}(f_1(X_{(i)}), f_2(X_{(i)})) \,.$$

In the second term, $\mathbb{E}(f_1(X_{(I)})\,|\,I)$ is the random variable equal to $\mathbb{E}(f_1(X_{(i)}))$ whenever $I = i$. Thus:

$$\begin{aligned}
&\operatorname{cov}(\mathbb{E}(f_1(X_{(I)})\,|\,I), \mathbb{E}(f_2(X_{(I)})\,|\,I)) \\
&= \frac{1}{n}\sum_{i=1}^{n} \mathbb{E}(f_1(X_{(i)}))\mathbb{E}(f_2(X_{(i)})) - \left(\frac{1}{n}\sum_{i=1}^{n} \mathbb{E}(f_1(X_{(i)}))\right)\left(\frac{1}{n}\sum_{i=1}^{n} \mathbb{E}(f_2(X_{(i)}))\right) \\
&= \frac{1}{2n^2}\sum_{i_1,i_2=1}^{n} \big(\mathbb{E}(f_1(X_{(i_1)})) - \mathbb{E}(f_1(X_{(i_2)}))\big)\big(\mathbb{E}(f_2(X_{(i_1)})) - \mathbb{E}(f_2(X_{(i_2)}))\big)\;.
\end{aligned}$$

The last equality comes from a classical manipulation, used in the proof of Chebyshev's sum inequality (see [S29, p. 94]). In fact, the whole argument here, can be seen as an application of Chebyshev's sum inequality. □

---


[S1] P. J. Gerrish, B. Galeota-Sprung, F. Cordero, P. Sniegowski, A. Colato, N. Hengartner, V. Vejalla, J. Chevallier, and B. Ycart, Natural selection and the advantage of recombination, Phys. Rev. Lett. **In Review** (2021).

[S2] P. J. Gerrish, F. Cordero, B. Galeota-Sprung, A. Colato, V. Vejalla, and P. Sniegowski, Natural selection promotes the evolution of recombination 2: during the selective process, Physical Review E **In Review** (2021).

[S3] P. J. Gerrish, B. Galeota-Sprung, P. Sniegowski, J. Chevallier, and B. Ycart, Natural selection promotes the evolution of recombination 1: among selected genotypes, Physical Review E **In Review** (2021).

[S4] W. J. Ewens, *Mathematical Population Genetics: I. Theoretical Introduction* (Springer, New York, NY, 2004).

[S5] N. Balakrishnan, Multivariate normal distribution and multivariate order statistics induced by ordering linear combinations, Stat. Probab. Lett. **17**, 343 (1993).

[S6] R. Song, S. G. Buchberger, and J. A. Deddens, Moments of variables summing to normal order statistics, Stat. Probab. Lett. **15**, 203 (1992).

[S7] J. B. Miller and A. Sanjurjo, A bridge from monty hall to the hot hand: The principle of restricted choice, J. Econ. Perspect. **33**, 144 (2019).

[S8] J. Berkson, Limitations of the application of fourfold table analysis to hospital data, Biometrics **2**, 47 (1946).

[S9] G. Gorjanc, P. Bijma, and J. M. Hickey, Reliability of pedigree-based and genomic evaluations in selected populations, Genet. Sel. Evol. **47**, 65 (2015).

[S10] E. M. Van Grevenhof, J. A. M. Van Arendonk, and P. Bijma, Response to genomic selection: the bulmer effect and the potential of genomic selection when the number of phenotypic records is limiting, Genet. Sel. Evol. **44**, 26 (2012).

[S11] G. M. Tallis, Ancestral covariance and the bulmer effect, Theor. Appl. Genet. **73**, 815 (1987).

[S12] M. Dupont-Nivet, J. Mallard, J. C. Bonnet, and J. M. Blanc, Evolution of genetic variability in a population of the edible snail, helix aspersa müller, undergoing domestication and short-term selection, Heredity **87**, 129 (2001).

[S13] M. G. Bulmer, Linkage disequilibrium and genetic variability, Genet. Res. **23**, 281 (1974).

[S14] M. G. Bulmer, The effect of selection on genetic variability, Am. Nat. **105**, 201 (1971).

[S15] K. Joag-Dev and F. Proschan, Negative association of random variables with applications, Ann. Statist. **11**, 286 (1983).

[S16] R. Szekli, *Stochastic ordering and dependence in applied probability*, L.N. Statist. No. 97 (Springer, New York, 1995).

[S17] D. Drouet Mari and S. Kotz, *Correlation and dependence* (Imperial College Press, London, 2001).

[S18] R. Pemantle, Towards a theory of negative dependence, J. Math. Phys. **41**, 1371 (2000).

[S19] J. Borcea, P. Bränden, and T. M. Liggett, Negative dependence and the geometry of polynomials, J. Amer. Math. Society **22**, 521 (2009).

[S20] T. Hu and J. Hu, Sufficient conditions for negative association of random variables, Statist. Probab. Letters **45**, 167 (1999).

[S21] T. Hu and J. Yang, Further developments on sufficient conditions for negative dependence of random variables, Statist. Probab. Letters **66**, 369 (2004).

[S22] M. Shaked and J. G. Shanthikumar, *Stochastic Orders* (Springer, New York, 2007).

[S23] B. Efron, Increasing properties of Pólya frequency function, Ann. Math. Statist. **36**, 272 (1965).

[S24] B.-E. Khaledi and S. Kochar, Stochastic comparisons and dependence among concomitants of order statistics, J. Multivariate Anal. **73**, 262 (2000).

[S25] H. A. David and H. N. Nagaraja, *Order statistics*, 3rd ed. (Wiley, New York, 2003).

[S26] T. Blessinger, More on stochastic comparisons and dependence among concomitants of order statistics, J. Multivariate Anal. **82**, 367 (2002).

[S27] I. Bayramoglu, B.-E. Khaledi, and M. Shaked, Stochastic comparisons of order statistics and their concomitants, J. Multivariate Anal. **124**, 105 (2014).

[S28] E. Amini-Seresht, B.-E. Khaledi, and M. Shaked, Order statistics with multivariate concomitants: stochastic comparisons, Statistics **50**, 190 (2016).

[S29] A. W. Marshall and I. Olkin, *Inequalities: theory of majorization and its applications* (Academic Press, New York, 1979).
